# Channeling macrophage polarization via selective translation inhibition by rocaglates increases macrophage resistance to Mycobacterium tuberculosis

**DOI:** 10.1101/691808

**Authors:** Sujoy Chatterjee, Shivraj M. Yabaji, Bidisha Bhattacharya, Emily Waligurski, Nandini Vallavoju, Somak Ray, Lauren E. Brown, Aaron B. Beeler, Alexander R. Ivanov, Lester Kobzik, John A. Porco, Igor Kramnik

**Affiliations:** Pulmonary Center, Department of Medicine, Boston University School of Medicine, National Emerging Infectious Diseases Laboratories (NEIDL), Boston University, Boston MA, 02118, USA; Department of Chemistry, Center for Molecular Discovery (BU-CMD), Boston University, Boston, MA, 02215, USA; Department of Environmental Health, Harvard School of Public Health, Boston, MA, 02115, USA; Barnett Institute of Chemical and Biological Analysis, Department of Chemistry and Chemical Biology, Northeastern University. Boston, MA, 02115, USA

**Author notes:** co-first authors (Sujoy Chatterjee, Shivraj M. Yabaji).

## Abstract

Macrophages contribute to host immunity and tissue homeostasis via alternative activation programs. M1-like macrophages control intracellular bacterial pathogens and tumor progression. In contrast, M2-like macrophages shape reparative microenvironments that can be conducive for pathogen survival or tumor growth. An imbalance of these macrophages phenotypes may perpetuate sites of chronic unresolved inflammation, such as infectious granulomas and solid tumors.

We have found that plant-derived and synthetic rocaglates sensitize macrophages to low concentrations of the M1-inducing cytokine IFN-gamma and inhibit their responsiveness to IL-4, a prototypical activator of the M2-like phenotype. Treatement of primary macrophages with rocaglates increased their resilience to oxidative stress, stimulated autophagy and killing of intracellular mycobacteria. Thus, rocaglates represent a novel class of immunomodulators that can direct macrophage polarization towards the M1-like phenotype in complex microenvironments associated with hypofunction of type 1 and/or hyperactivation of type 2 immunity, e.g. chronic bacterial infections, allergies and, possibly, certain tumors.

## Introduction

The increasing problem of infectious diseases caused by antibiotic-resistant bacteria has intensified a search for novel therapeutic approaches. Among them are host-directed therapies (HDT) aimed at either boosting immune-mediated bacterial control or reducing immunopathology, broadly referred to as mechanisms of host resistance or disease tolerance, respectively (1). Many intracellular bacterial pathogens reside in macrophages, the very cells of innate immune system whose major function is to eliminate invading pathogens. This paradox is enabled by macrophages’ plasticity, which successful pathogens exploit to create cellular niches for persistence and replication within inflamed tissue of susceptible hosts.

To combat intracellular bacteria, macrophages activate cell autonomous defense mechanisms, such as phagocytosis, production of highly toxic reactive oxygen and nitrogen species, bactericidal peptides, as well as phagosome maturation and autophagy to deliver the ingested pathogens to lysosomes for destruction. This pro-inflammatory type of macrophage activation can be induced by microbial ligands and proinflammatory cytokines. It is broadly referred to as “classical” or M1 type, although this definition encompasses an array of related, but non-identical macrophage activation states. Individuals whose macrophages fail to respond to IFNγ, the prototypical M1 polarizing signal, are extremely susceptible to infections caused by intracellular mycobacterial species, including an avirulent vaccine strain of *M.bovis* BCG and a normally avirulent environmental non-tuberculous mycobacteria (NTM). The alternative macrophage activation programs, broadly referred to as M2 type, are mediated by upregulation of anti-inflammatory cytokines and characterized by an increased macrophage permissiveness for intracellular bacteria. Activation of an alternative macrophage activation program, M2 type, by IL-4 favors *M.tb* replication by suppressing autophagy and antigen presentation by the infected macrophages. Overexpression of IL-4 in a mouse model in vivo leads to progression of tuberculosis (TB) and formation of organized necrotic granulomas typical of human TB(2). The M2-like differentiation of macrophages is also involved in local immune suppression within tumor microenvironments and tumor promotion (3). However, the M2-like macrophages play important homeostatic and adaptive physiological roles in immune regulation, tissue remodeling, fibrosis and control of parasitic helminth infections(4). Within specific in vivo environments macrophages may be exposed to multiple and often conflicting polarization signals. For example, a helminth-induced type 2 immune response antagonizes host protective responses to mycobacteria(5).

In vitro, the prototypic M1- and M2-like macrophage polarization states can be artificially induced by treatment with cytokines or microbial ligands and represent polar antagonistic phenotypes. Thus, pretreatment with IL-4 was shown to downregulate the macrophage responsiveness to IFNγ (5–9), while priming with IFNγ reduces their responses to IL-4(10). In vivo, macrophage polarization is influenced not only by systemic cytokine levels, but also by the composition and anatomy of specific lesions, such as TB granulomas or tumors that serve as local sources of polarizing cytokines and growth factors. Despite the mutual antagonism of these macrophage polarization programs, recent studies demonstrated simultaneous presence of M1- and M2-like macrophages within TB lesions(11, 12). This suggests that the balance of M1/M2 macrophage phenotypes locally within TB lesions may determine trajectories of the individual granuloma progression(13). Spatial transcriptomic analysis and immunochemistry revealed that the M2-like macrophage markers were more abundant within the inner areas of TB granulomas (11, 12), whereas the IFNγ-producing T cells were located mostly on the periphery of the organized granulomas(14). Because IFNγ is a labile homodimer, the diffusion of its biologically active form within inflammatory lesions is limited. Therefore, we searched for small molecules that enhance M1-like macrophage polarization in synergy with low concentrations of IFNγ capable of directing granuloma macrophages towards the M1-like activation and enhancing their anti-mycobacterial activity.

Screening small molecule libraries, we found several synthetic rocaglates that potentiated pleiotropic effects of IFNγ on primary macrophages (15). Rocaglates are a class of bioactive derivatives of natural products isolated from medicinal plants of the *Aglaia* species that have been used in traditional Chinese medicine for treatment of fever, cough, diarrhea and inflammation. In modern medicine, the therapeutic potential of synthetic rocaglates has been evaluated for cancer therapy (16–19) and viral infections including coronaviruses(20). Rocaglates bind to and suppress the RNA helicase activity of the eIF-4A subunit of the eIF4F translation initiation complex and selectively inhibit cap-dependent protein translation (16, 21). Also, the structurally related rocaglamide A (RocA) has been shown to inhibit the Raf-MEK-ERK growth factor activated pathway by targeting prohibitins 1 and 2(22).

Although rocaglates are known as potent translation inhibitors, we found that in the presence of IFNγ at concentrations as low as 0.1 U/ml, select rocaglates induced the upregulation of IRF1 protein, a master regulator of IFNγ-activated pathways. In the present study, we delineate mechanisms of the enhancement the M1 macrophage polarization by active rocaglates, demonstrate that rocaglates also inhibit macrophage responsiveness to CSF-1 and IL-4 and potentiate killing of virulent Mtb by primary macrophages.

## Results

### Macrophage activation by rocaglates is coupled to partial translational inhibition

To delineate the relationship between translation inhibition and macrophage activation by rocaglates, we screened a library of synthetic rocaglate derivatives. Out of 86 compounds, twenty-three induced significant *Irf1* mRNA upregulation in the presence of 0.2 U/ml of IFNγ. Next, we determined the translation inhibitory concentrations IC_50_ of these compounds using the 293TR-FLuc translation reporter cell line (16). We observed a tight negative correlation (r = - 0.6832) between the *Irf1* mRNA induction and IC_50_ of those compounds (**Fig.1A, Suppl. Table 1**) demonstrating mechanistic link between the translational inhibition and *Irf1* co-stimulation properties. To verify this relationship in primary macrophages, first we compared translation inhibition activities of two structurally similar rocaglates CMLD010536 and CMLD010850 (**Fig.1B**) in mouse bone marrow-derived macrophages (BMDM) using a puromycin incorporation assay (**Fig 1 C**)(23). CMLD010536 exerted greater translation inhibition than CMLD010850 in primary macrophages. Next, we compared the induction of target genes by these compounds (**Fig.1D**): CMLD010536 was significantly more potent as an inducer of *Irf1* mRNA expression cooperatively with low dose of IFNγ, as well as of another, IFNγ - independent, rocaglate target gene *Txnip* (16).

**Figure 1.**
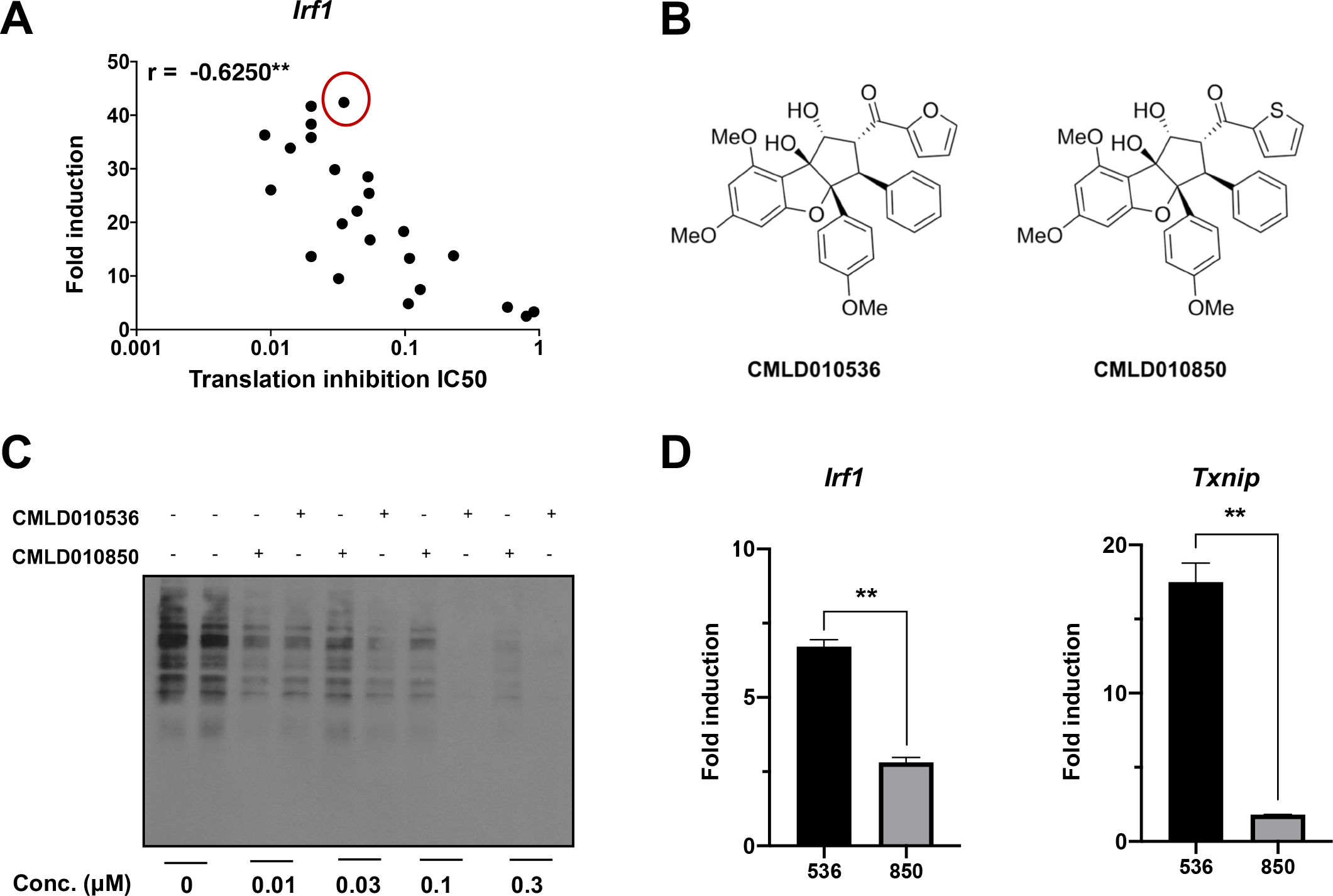
Translation inhibition is required for rocaglate to activate BMDM. **(A)** Twenty three rocaglate analogues were examined (at 0.3 μM for 24 h) for their translation inhibition activity and induction of *Irf1* mRNA expression in the presence of 0.1 U/ml of IFNγ. The correlation coefficient was calculated using GraphPad Prism 8. **(B)** Chemical structure of small molecules CMLD010536 and CMLD010850. **(C)** Translation inhibition by rocaglates in BMDMs assessed using puromycin incorporation assay. BMDMs were treated with indicated concentrations of CMLD010536 and CMLD010850 for 2 h followed by puromycin (5 μg/ml) for 1 h. Puromycin incorporation was monitored by Western blot using puromycin-specific antibody. **(D)** BMDMs were treated with 100 nM CMLD010536 (labeled as 536) and CMLD010850 (labeled as 850). The *Irf1* (in the presence of IFNγ) and *Txnip* (without IFNγ) mRNAs induction was assessed using quantitative real time RT-PCR (qRT-PCR). The data are represented as mean ± SEM and *p value* ≤0.05 was considered statistically significant.

Taken together these data demonstrate that macrophage activation by rocaglates is proportional to their translation inhibition activity. Importantly, general translational inhibitors, such as cycloheximide and puromycin, do not cause IRF1 co-stimulation, as described in our previous studies (15).

### Activation of the p38 stress kinase and the integrated stress response pathways mediate the macrophage activation by rocaglates

To further investigate the dynamics of translation inhibition by CMLD010536 in BMDM, we used a puromycin incorporation assay. Puromycin incorporation during a one-hour pulse was determined after 3, 9 and 24 h of rocaglate treatment using western blot with puromycin-specific antibodies. The rocaglate treatment significantly reduced translation rates at all timepoints in a dose-dependent manner (**Fig.2A**). However, the levels of ongoing translation were similar at all of the timepoints demonstrating that translation inhibition was partial and sustained over the time course of the rocaglate treatment. To identify proteins expressed in BMDMs after 24 h of treatment with 100 nM of CMLD010536, we performed proteomic analysis using mass spectrometry (**Suppl. Table 2**). The abundance of Sequestosome 1 (SQSTM1, aka p62) protein was found to significantly increase. We confirmed the induction of p62 protein and mRNA (**Fig.2 B** and 2**C**), demonstrating that treatment of BMDM with the rocaglate induced de novo p62 protein and mRNA syntheses.

**Figure 2.**
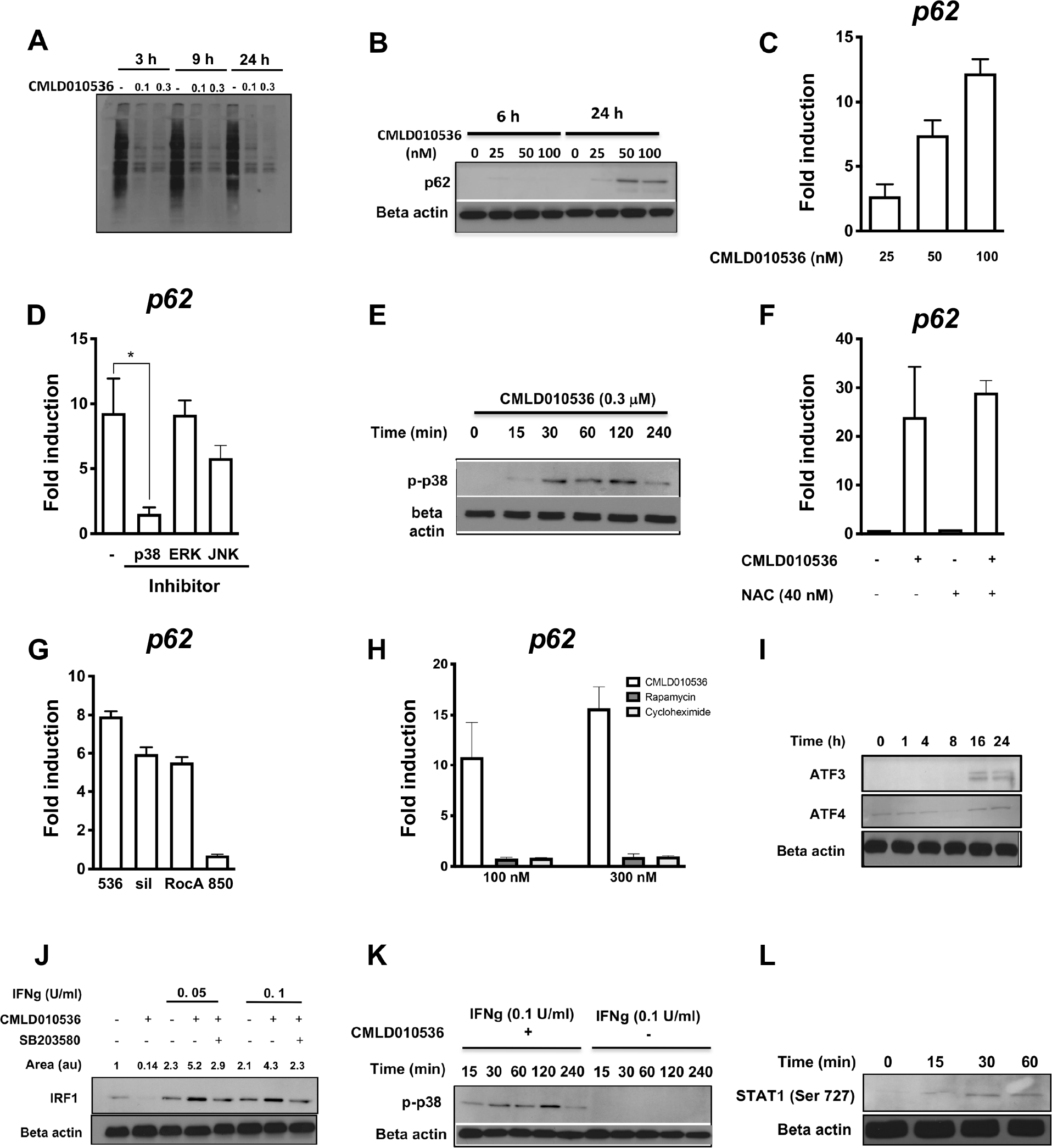
Selective translation inhibition by rocaglate remodels macrophage proteome and upregulates stress response proteins. **(A)** BMDMs were treated with 0.1 and 0.3 μM CMLD010536 for 2, 8 and 23 h followed by a treatment with puromycin (5 μg/ml) for additional 1 h. Incorporation of puromycin was monitored by Western blot using puromycin-specific antibody. **(B)** BMDMs were treated with CMLD010536 (0, 25, 50 and 100 nM) for 6 and 24 h. The expression of p62 protein was measured by Western blot using p62-specific antibody. **(C)** The *p62* mRNA induction in BMDMs treated with CMLD010536 (at 25, 50 and 100 nM) for 6 h. The *p62* mRNA levels were determined using qRT-PCR and expressed as a fold change compared to untreated cells. **(D)** Effect of MAPK inhibitors SB203580 (p38), SB600125 (JNK) andU0126 (ERK) on *p62* mRNA expression in BMDM treated with 100 nM CMLD010536 for 6 h. Inhibitors were added to a final concentration 10 μM, 30 min prior to the rocaglate treatment. The *p62* mRNA levels were determined using qRT-PCR and expressed as fold induction compared to untreated BMDMs. **(E)** The kinetics of p38 phophorylation in BMDMs treated with 0.3 μM CMLD010536 was determined by Western blot using phospho-p38 specific antibody. **(F)** The expression of *p62* mRNA induced by CMLD010536 (100 nM for 6 h) was not affected in the presence of ROS scavenger NAC (40 nM). The mRNA levels were determined using qRT-PCR. NAC was added 1 h before the rocaglate treatment. **(G)** Effect of the rocaglate derivatives CMLD010536 (536), sivestrol (sil), rocaglamide A (RocA) and CMLD010850 (850) on *p62* mRNA expression. BMDMs were treated with each compound at 100 nM for 24 h. The *p62* mRNA levels were measured by qRT-PCR and expressed as the fold change compared to untreated cells. **(H)** Effects of CMLD010536, rapamycin and cycloheximide added to final concentrations 100 and 300 nM for 6 h on the *p62* mRNA expression. The p62 levels were determined as in G. **(I)** The kinetics of the ATF3 and ATF4 protein levels in BMDMs treated with 100 nM CMLD010536 as determined by Western blot. **(J)** The effect of the p38 inhibitor SB203580 (10 μM) on IRF1 protein induction by CMLD010536 (300 nM) in synergy with low concentrations of IFNγ (0.05 or 0.1 U/ml). The inhibitor was added to BMDMs 30 min prior to stimulation with the rocaglate and IFNg. The IRF1 protein levels were determined after 6 h by Western blot using IRF1-specific antibody. The relative densitometric values of IRF1 bands’ areas were determined after normalization and are shown as ratio to non-stimulated controls above the blot. **(K)** Treatment with 0.1 U/ml alone did not induce the p38 phosphorylation in BMDMs as compared to cells treated with 0.1 U.ml of IFNγ in combination with CMLD010536 (0.3 μM). BMDMs were treated with 0.1 U/ml of IFNγ either in the absence or presence of the rocaglate and phospho-p38 levels were determined by Western blot using phospho-p38 specific antibody. **(L)** BMDMs were treated with 0.3 μM CMLD010536 and phosphorylation of STAT1 at Ser727 at indicated timepoints was monitored by Western blot using phospho-Ser727-specific antibody.

SQSTM1/p62 plays an important role in stress resilience by controlling selective autophagy and anti-oxidant defense gene expression (24–33). Its expression can be induced by various stressors. Our previous analysis of CMLD010536-activated pathways using gene expression profiling suggested activation of stress kinase-mediated pathways (15). Therefore, we tested whether p62 induction by CMLD010536 is mediated by stress-activated MAP kinases. The p38 inhibitor (SB203580 at 10 uM), added 30 min prior to treatment with 100 nM CMLD010536, completely abrogated *p62* mRNA upregulation. The effect of the JNK inhibitor (SB600125) was less prominent, while ERK inhibition (U0126/PD98059) had no effect (**Fig.2D**). The p38 MAPK inhibitor also abrogated the expression of another rocaglate-induced inflammatory gene, *Txnip* (**Suppl. Fig.1A**). The MAP kinase inhibitors displayed similar effects on the p62 protein levels (**Suppl.Fig.1B**). We also observed a dramatic downregulation of another rocaglate-inducible mRNA, *Ptgs2*, by the p38 inhibitor (**Suppl.Fig.1C**). Next, we compared the kinetics of p38 phosphorylation (**Fig.2E**) and *p62* mRNA upregulation (**Suppl.Fig. 2A**) during the course of the rocaglate treatment: rapid activation of p38, started at 15 min of the treatment, reached a plateau at 30 – 120 min and declined by 4 h. Meanwhile, the *p62* mRNA continued to increase 4 – 24 h of the treatment duration indicating that p38 phosphorylation was necessary to initiate a signaling cascade leading to the p62 upregulation, but was dispensable for its maintenance. Because p38 activation can be induced by reactive oxygen species (ROS), we tested whether oxidative stress plays a role in the p38-mediated p62 induction by the rocaglate. However, *Sqstm1/p62* mRNA induction was unaffected in the presence of ROS scavenger N-acetyl-cysteine (NAC) (**Fig.2F**). Comparing the *p62*-inducing activities of CMLD010536 with natural rocaglates silvestrol and rocaglamide A demonstrated that those compounds had similar activities, while CMLD010850 did not induce *p62* mRNA expression (**Fig.2G**). Notably, another suppressor of cap-dependent translation, rapamycin (which is known to inhibit translation via inhibition of mTOR pathway), as well as the known translation elongation inhibitor cycloheximide, both failed to stimulate the expression of *p62* (**Fig.2H**).

Cap-dependent protein translation can be inhibited by several stress-activated kinases via phosphorylation of the translation initiation factor eIF2α, thus initiating a signaling cascade known as the Integrated Stress Response (ISR). Inhibition of *cap-dependent* translation, initiates a *cap-independent* translation of a stress response transcription factor ATF4 followed by the upregulation of its downstream target ATF3. We observed that both ATF4 and ATF3 proteins (**Fig.2I)** and ATF3 mRNA (**Suppl.Fig.2B**) were upregulated at 16 – 24 h of the CMLD010536 treatment.

Taken together, these data demonstrate that macrophage treatment with rocaglates induces step-wise transcriptome and proteome remodeling. At the early initiation step (0.5 – 2 h), selective translation inhibition is associated with the activation of p38 stress kinase that drives transcriptional upregulation of *p62*, *Ptgs2, Txnip* and other inflammatory genes. The translation of ISR transcription factors and *p62* occurs with a delay at a later stage (16 – 24 h) of rocaglate treatment. At this later stage, the activating effects of rocaglates on macrophages mimic diverse stressors whose activities converge on a common stress response module – ISR. These effects are consistent with the known ability of rocaglates to inhibit eIF4A helicase, a component of the cap-dependent translation initiation complex. In contrast to other translation inhibitors, such as cyclohexamide and mTOR inhibitors, the effect of rocaglates is partial and more selective, allowing for the upregulation of proteins that activate stress adaptation pathways.

### The p38 activation mediates synergy of rocaglates with low concentrations of IFNγ

In the presence of low concentrations of IFNγ, CMLD010536 upregulates mRNA and protein expression of *Irf1*, a key transcription factor in the IFNγ network. The upregulation of the *Irf1* gene expression by IFNγ is mediated by the binding of STAT1 homodimers to the gamma-activatable sequence (GAS) in the *Irf1* promoter. The STAT1 homodimer formation and promoter binding require STAT1 phosphorylation at the canonical tyrosine-701 by JAK kinases, an event that occurs upon the IFNγ dimer binding to its receptor (34, 35). However, CMLD010536 did not increase STAT1 phosphorylation at the canonical tyrosine site in our studies. Therefore, we tested whether the synergistic effect of the rocaglate on IRF1 induction was mediated via alternative mechanism(s) involving MAP kinase activation. There is an emerging evidence that serine (Ser-727) phosphorylation of STAT1 by stress-activated MAP kinases p38 and JNK mediates various stress responses (36) and may account for nearly 80% of IFNγ-induced transcriptional activity. Indeed, CMLD010536 boosted the levels of IRF1 protein in the presence of IFNγ at concentrations as low as 0.05 and 0.1 U/ml, and this boosting effect was abrogated by p38 inhibition (**Fig.2J)**. The p38 inhibitor also suppressed the *Irf1* mRNA co-stimulation by the rocaglate (**Suppl.Fig.3A**). In contrast to the rocaglate treatment, low concentrations of IFNγ alone did not induce the p38 phosphorylation (**Fig.2K**). Finally, we found that the CMLD010536 treatment induced STAT1 Ser727 phosphorylation as early as 15 min, which was maintained for at least 4 hours (**Fig.2L**).

These data demonstrate that the p38 MAPK pathway activation by rocaglates plays a central role in their synergy with IFNγ in macrophage activation. Importantly, this STAT1 modification does not increase DNA binding or nuclear translocation of STAT1 dimers (37–39). Indeed, treatment with active rocaglates alone induced no *Irf1* mRNA upregulation(15) and even suppressed the basal IRF1 protein expression (**Fig.2J**). Furthermore, the p38 inhibition did not suppress the *Irf1* mRNA induction by standard doses of IFNγ alone (**Suppl.Fig.3B**). Thus, the effect of rocaglates does not simply mimic the canonical IFNγ pathway. We propose that low doses of IFNγ are necessary to initiate the STAT1 dimer formation and binding to the IRF1 promoter, while the p38 activation by rocaglates and STAT1 Ser-727 phosphorylation boosts and sustains the *Irf1* transcription, most likely, by facilitiating interactions of STAT1 with transcriptional co-activators.

### Suppression of the alternative activation of macrophages by rocaglates

Once we determined that CMLD010536 boosted a pro-inflammatory (M1-like) macrophage program, we investigated whether it could concurrently suppress the alternative (M2-like) macrophage activation. In agreement with the known antagonistic relationship of these modes of macrophage activation, pretreatment of macrophages with IFNγ abrogated their response to the M2-polarizing cytokine IL-4, as measured by the *Arg1* mRNA expression (**Fig.3A**). Similarly, the induction of the M2 polarization markers *Arg1* and *Fizz1* by IL-4 was suppressed by CMLD010536 in a dose-dependent manner (**Fig.3B and Suppl.Fig.4A,** respectively). Interestingly, this suppressor effect of the rocaglate does not require the co-stimulation with IFNγ. It does correlate, however, with a rocaglate ability to inhibit protein translation: at 0.1 mM concentration the alternative macrophage activation was suppressed by CMLD010536, silvestrol and rocaglamide A, but not by CMLD010850 (**Fig.3C**), which is a much less potent translational inhibitor and does not inhibit translation at this concentration (**Fig 1C**).

**Figure 3.**
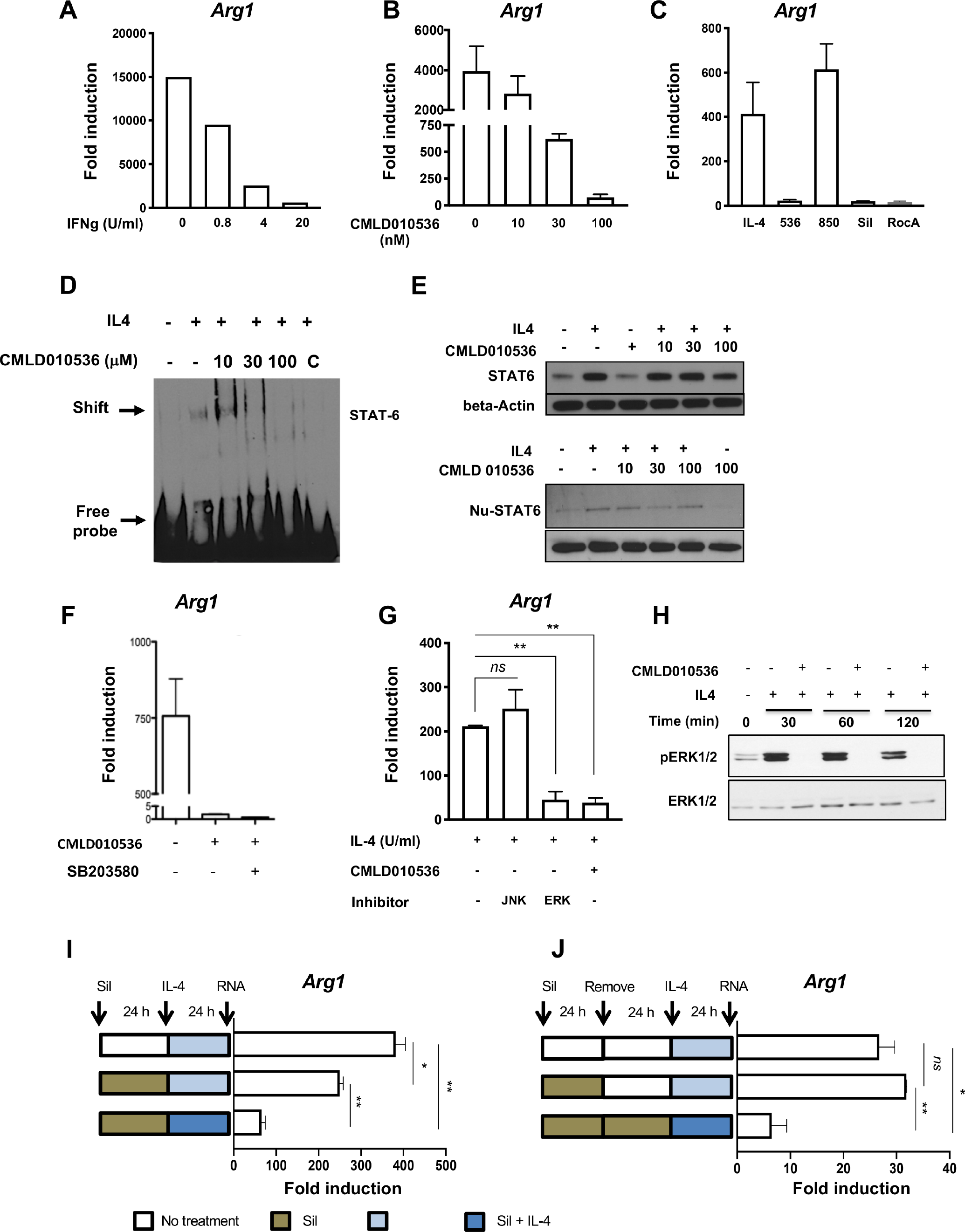
Suppression of the alternative macrophage activation by rocaglates. **(A)** Pretreatment with IFNg suppresses macrophage responses to IL-4. BMDMs were pretreated with increasing concentrations of IFNγ for 4h and stimulated with 25 ng/ml IL4 for additional 20 h. The expression of *Arg1* mRNA was measured using qRT-PCR. Fold induction was calculated using the IL4 untreated controls. **(B)** BMDMs were pretreated with indicated concentrations of CMLD010536 for 4h, stimulated with IL4 and analyzed for *Arg1* mRNA expression as in A. **(C)** BMDMs were pretreated with different rocaglate derivatives (at 100nM) for 4 h followed by treatment with 25 ng/ml IL-4 for 20 h. The *Arg1* mRNA induction was determined as in A. **(D**) Effect of CMLD010536 on the IL4-induced STAT6 binding activity in BMDMs was monitored using EMSA. BMDMs were pretreated with the indicated rocaglate concentrations for 2 h and stimulated with IL4 (25 ng/ml) for additional 4 h. Nuclear extracts were prepared and STAT6 promoter binding activity was monitored by EMSA using STAT6 specific DNA probe. Competition with unlabelled STAT6 probe (C) demonstrates the specificity of the shifted band. **(E)** Rocaglate do not inhibit STAT6 protein upregulation by IL4. BMDMs were treated with indicated concentrations of CMLD010536 for 2 h followed by a treatment with IL4 (25 ng/ml) for additional 4 h. Total (upper panel) and nuclear (lower panel) STAT6 protein levels were measured by Western blot. The total STAT6 protein levels were not affected by the rocaglate treatments, while the nuclear STAT6 levels appear to be reduced. **(F)** The induction of the *Arg1* mRNA by IL4 (25 ng/ml) was suppressed by CMLD010536 (100 nM) and this suppression was not abbrogated by the p38 inhibitor SB203580 (10 μM). The *Arg1* mRNA was measured using qRT-PCR at 24 h. **(G)** The induction of the *Arg1* mRNA by IL4 (25 ng/ml) was not affected by the JNK inhibitor (SB600125), but was suppressed by the ERK inhibitor (U0126) to a degree similar to CMLD010536 (100 nM). The inhibitors were added 30 min prior to IL-4 and the *Arg1* mRNA expression was measured using qRT-PCR at 24 h. **(H)** The rocaglate treatment abrogates ERK1/2 phosphorylation in IL-4 stimulated macrophages. BMDMs were pre-treated with CMLD010536 (100 nM) for 12 h and stimulated with IL4 (100 ng/ml). Phospho and total ERK1/2 proteins were measured using Western blot after the indicated periods of the IL-4 stimulation. **(I and J)** The suppression of the BMDM response to IL-4 by the rocaglate is reversible. BMDMs were pretreated with CMLD010536 (100 nM) for 24 h, washed and stimulated with IL-4 (100 ng/ml) either alone or in the presence of CMLD010536 (100 nM). The cells not treated with CMLD010536 and stimulated with IL-4 were used as a positive control. The expression of *Arg1* mRNA was measured by using qRT-PCR as above. In (J) BMDMs were pretreated and washed as in I, but were rested for additional 24 h before stimulation with IL-4. BMDMs cultured in the presence of the rocaglate for the duration of the experiment served as a control for suppression. The data are represented as mean ± SEM and *p values* ≤0.05 were considered significant.

The upregulation of the IL-4-inducible genes is mediated by the transcription factor STAT6. Using electrophoretic mobility shift assay (EMSA) with a STAT6 consensus sequence oligonucleotide probe, we determined that pretreatment with 30 and 100 nM of CMLD010536 prior to IL-4 stimulation inhibited STAT6 DNA binding (**Fig.3D**). This effect was not associated with a decrease in the total STAT6 protein levels (**Fig.3E**, upper panel). However, nuclear STAT6 protein was somewhat reduced (Fig.3E, lower panel). These observations suggested that the suppression of the STAT6 DNA binding activity was a regulatory phenomenon that could not be simply attributed to inhibition of the STAT6 protein translation. Therefore, we sought to determine whether the suppressor activity was mediated by MAP kinases. The p38 inhibitor, however, did not restore the expression of *Arg1* inhibited by CMLD10536 (**Fig.3F**). Instead, we observed that the rocaglate treatment mimicked the inhibitory effect of the ERK inhibitor U0126 on IL4-induced *Arg1* mRNA expression (**Fig.3G**). Indeed, the rocaglate treatment completely abrogated the ERK1/2 phosphorylation in IL4-stimulated macrophages without affecting the total ERK1/2 levels (**Fig.3H**). Although it remains to be determined whether rocaglates inhibit ERK activation directly, or block upstream activators of ERK, this finding is consistent with the previously described role of ERK1/2 in the alternative macrophage activation. One of the relevant direct rocaglate targets in this pathway could be Myc, whose activity is necessary for M2-like macrophage polarization (40, 41) and whose translation was shown to be selectively inhibited by rocaglates(42).

Next, we wanted to determine whether the rocaglates treatment induces transient or stable suppression of the M2-like phenotype. As shown in **Fig.3I**, pre-treatement and removal of the rocaglate prior to the IL-4 stimulation had a substantially weaker inhibitory effect on the *Arg1* gene expression, as compared to simultaneous treatment with the rocaglate and IL-4. When we increased the interval between the rocaglate treatment and IL-4 stimulation to 24 h, the inhibitory effect of the rocaglate pretreatment disappeared completely (**Fig.3J**). Similar effects were observed when *Fizz1* gene expression was used as an M2 polarization marker (**Suppl.Fig.4B and 4C**). Thus, the rocaglate inhibition of IL-4 responses is transient and can be completely reversed after their removal.

These experiments demonstrated that, unlike the rocaglates’ effects on the pro-inflammatory gene expression, the inhibition of alternative macrophage activation by rocaglates is driven by a separate, p38-independent, pathway. Selective translation inhibition by rocaglates, however, is required for the activation of both pathways. Perhaps, they represent two modules of a coordinated physiological response of macrophages to attenuated protein translation – a common sign of, and an adaptation to, ongoing stress.

### Macrophage activation by rocaglates improves bacterial control

The ability of rocaglates to synergize with low concentrations of IFNγ, inhibit the alternative macrophage activation and activate protective stress responses, make this class of compounds an attractive candidate for host-directed therapy of infections caused by intracellular bacterial pathogens. The SQSTM1/p62 protein, which is induced by rocaglates in an p38-dependent, but IFNγ-independent manner, coordinates activation of central stress response pathways: antioxidative defense and autophagy (24, 25, 28–33, 43). It is a major adaptor protein that targets intracellular bacteria to LC3-positive autophagosomes(44). Previously, we have reported that the rocaglate treatment stimulates autophagic flux and increases the lipidated form of LC3 associated with autophagosome maturation (15). To test whether rocaglate treatment could enhance macrophage resistance to intracellular bacteria, we treated BMDM with CMLD010536 (50 nM), infected them with *Mycobacterium bovis* BCG, and determined their bacterial loads and viability at 24 h p.i.. The rocaglate treatment significantly reduced the bacterial loads, as determined by quantitative PCR of the bacterial genomes (**Fig.4A**), and viability, as determined by CFU plating (**Suppl. Fig.5A**). Using fluorescent *M. bovis* BCG (BCG:*gfp*) and staining with Lysotracker, we visualized the delivery of the invading bacteria to lysosomes at 1, 24 and 48 h post infection. Indeed, pretreatment with CMLD010536 significantly increased the BCG:*gfp* colocalization with lysosome at 24 and 48 h (**Fig.4B** and **4C**). These data demonstrate that rocaglate-mediated macrophage activation significantly increased clearance of a vacuolar intracellular bacteria (BCG).

**Figure 4.**
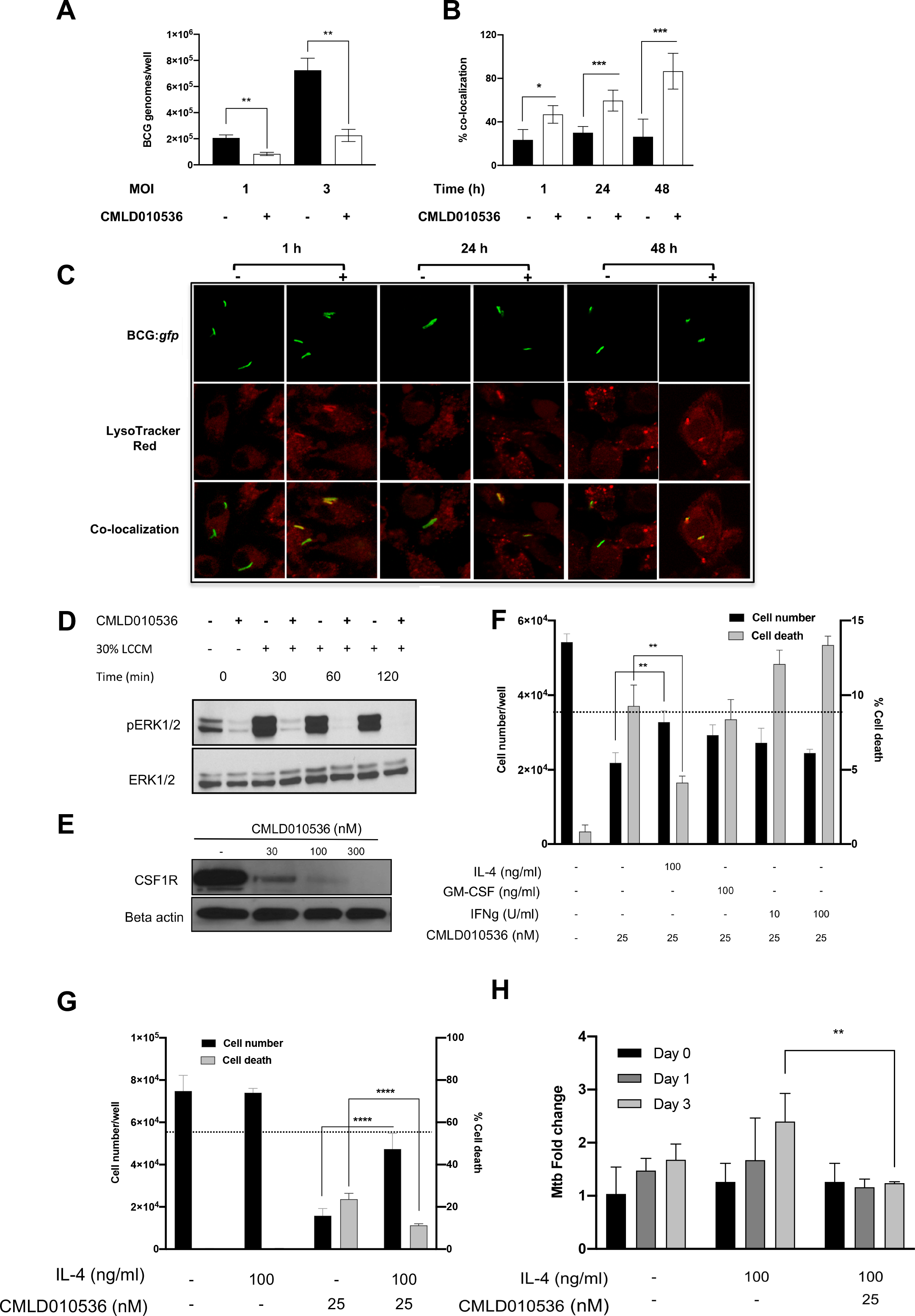
**Rocaglates stimulate antimycobacterial defenses** in BMDMs. (**A**) BMDMs were pretreated with CMLD010536 (50 nM) for 24 h and subsequently infected with *M. bovis* BCG at MOI 1 and 3. The intracellular BCG loads were measured 24 h post infection by determining genome equivalents using qPCR with BCG-specific primers and Taqman probe. (**B and C**) Rocaglates promote phagosome-lysosome fusion. BMDMs were pretreated with 25 nM CMLD010536 for 6 h and infected with *M. bovis* BCG expressing *gfp* (BCG:*gfp*) at MOI 1. At 1, 24 and 48 h p.i. the cells were strained with Lysotracker Red for 1 h, processed for imaging by confocal microscopy and analyzed for co-localization of lysosomes with phagosomes containing BCG:*gfp*. The percentage of BCG-positive phagosomes co-localization wth lysosomes and representative images are presented in panels B and C, respectively. The data in B are represented as mean + SEM (n=50, phagosomes), p<0.05. (**D and E**) Rocaglate inhibit ERK phosphorylation induced by CSF1-containing media (LCCM)(D) and the expression of CSF1R (E). BMDMs were pre-treated with 100 nM CMLD010536 for 12 h and subsequently treated with 30% LCCM for the indicated periods of time. (**D**) Phospho- and total ERK1/2 protein levels were determined by Western blot. **(E)** BMDMs were treated with indicated concentrations of CMLD010536 for 24 h and CSF1R protein levels were determined by Western blot. (**F**) BMDMs were treated with 25 nM CMLD010536 either alone or in combination with IL-4 (100 ng/mL), GM-CSF (100 ng/mL) or IFNg (10 and 100 U/mL) for 72 h. Total cell numbers (left Y axis) and percentage cell death (right Y axis) were analysed using LIVE/DEAD STAINIG and automated micropscopy. The dotted line indicates the cell number at day 0. **(G)**. IL-4 improves the survival of the rocaglate treated and Mtb-infected macrophages. BMDMs were pretreated with IL-4 (100 ng/mL) for 16 h and infected with Mtb H37Rv at MOI 1. CMLD010536 (25 nM) was added 2 h p.i. either alone or in combination with IL-4 (100 ng/mL). Total cell numbers and percent of dead cells at 72h p.i. were determined as in F. The dotted line denotes the cell numbers immediately after the Mtb infection and washing off the extracellular bacteria (day 0). **(H)** The rocaglate treatment reduces the Mtb load in the presence of IL-4. The intracellular bacterial loads were determined after 2 h phagocytosis (Day 0), and at days 1 and 3 p.i. using qPCR and normalized using BCG spike as an internal control. The Mtb loads are presented as fold change compared to Mtb uptake by untreated BMDMs at 0h. The data are represented as mean ± SEM; *p value* ≤0.05 was considered statistically significant.

Compared to avirulent vaccine BCG, virulent *Mycobacterium tuberculosis* (Mtb) is more resilient to macrophage attack. In naïve macrophages, the bacteria block phagosome – lysosome fusion, then escape from phagosomes to the cytoplasm, grow and eventually destroy the host cells. Because of a slow replication rate of the bacteria this cycle takes several days, unless a high MOI is used. To test whether rocaglates would improve the macrophage ability to control virulent Mtb in vitro, we used low multiplicity of infection (MOI) and extended the time post infection (p.i.) to 72 h. However, the viability of BMDMs infected with Mtb and treated with CMLD10536 at 25 – 50 mM for 3 days was low. Even in the absence of infection, the rocaglates treatment increased cell death by that time (**Suppl.Fig.5B**). The BMDMs population prepared in our laboratory is critically dependent on CSF1 for survival (45). Nevertheless, increasing the CSF1-containing conditioned media 3-fold did not improve cell survival in the presence of rocaglates. We also observed that ERK1/2 phosphorylation, which is downstream of CSF1 receptor (CSF1R) signaling, could not be restored in the rocaglates-treated cells by increasing CSF1 (**Fig.4D**). Finally, we found that rocaglates suppressed the expression of CSF1 receptor (**Fig.4E**). Therefore, we attempted to identify alternative survival factors and compared the survival of the rocaglates-treated BMDMs in the presence of GM-CSF, IFNγ and IL-4 (**Fig.4F**). Surprisingly, IL-4 significantly improved the the macrophage survival within 72 h, as compared to GM-CSF and IFNγ. However, IL-4 did not restore macrophage proliferation in the presence of the rocaglate (**Suppl.Fig.5D**). The pro-survival effect of IL-4 on rocaglate-treated BMDMs was not mediated by either STAT6 or ERK1/2, since both were inhibited by rocaglate in the presence of IL-4 (**Fig.3D and 3H**). Possibly, this effect is mediated by activation of the Akt pathway downstream of the IL-4 receptor(46).

The pro-survival effect of IL-4 was also observed in Mtb-infected BMDM cultures at 72 h p.i.(**Fig.4G**). In contrast, GM-CSF failed to improve the macrophage survival in these settings (**Suppl.Fig.5C**). Therefore, we tested the effects of rocaglates on Mtb replication in macrophages in the presence of IL-4. The bacterial loads were estimated using an optimized quantitative PCR method developed in our laboratory (Yabaji et al., submitted for publication). We observed an approximately 2-fold increase between 0 – 72 h p.i. in naïve macrophages (**Fig.4H**). Pretreatement with IL4 had no effect on the bacterial uptake. In the presence of IL-4, the bacterial loads increased by 72 h p.i, as compared to naïve macrophages. In contrast, no bacterial growth was observed in BMDMs treated with rocaglate and IL-4, and the bacterial loads at 72 h p.i. were significantly lower as compared to naïve and IL-4-treated macrophages. Thus, rocaglate improves macrophage ability to control Mtb growth even in the presence of a M2-polarizing cytokine. Notably, ARG1-positive (M2-polarized) macrophages have been detected in TB granulomas in mice, non-human primates and humans (47). Therefore, this rocaglate activity may be beneficial in the context of active disease.

To begin testing effects of rocaglates in vivo, we have chosen a model of acute respiratory challenge with *Streptococcus pneumoniae* (serotype 3)(48). Alveolar macrophages (AM) play an important role at the early phase of this infection (49, 50). It is also worth noting that in naïve hosts the AM displays an M2-like phenotype. We wondered whether rocaglate pretreatment could prime macrophages for more efficient control of the acute bacterial infection. Outbred CD-1 IGS male mice were treated with CMLD010536 (25 g/kg, i.p.) at 48, 24 and 0 hours before the respiratory challenge with *Strep. pneumoniae*. At 24 h p.i., bronchoalveolar lavage (BAL) was performed to harvest 2 ml of BAL fluid. We observed significant reduction of live bacteria (as determined by BAL CFU counts, **Suppl.Fig.6** left panel) and BAL cell numbers (**Suppl.Fig.6**, right panel) in rocaglate pretreated mice as compared to vehicle-treated controls. These results show that rocaglate pretreatment increased bacterial control and reduced inflammatory cell recruitment to the lungs at the early stage of respiratory infection with *Strep. pneumoniae*.

## Discussion

Taken together, our data demonstrate that rocaglates mimic pleiotropic effects of IFNγ on primary macrophages. These effects are dependent on their translation inhibition activity and are partially mediated by the activation of p38 MAPK, p62 and integrated stress response modules. In contrast, they cause suppression of alternative macrophage activation and growth factor responses mediated by STAT6, CSF-1 receptor and ERK. Thus, exposure of primary macrophage to rocaglates restricts their differentiation, channeling it towards the M1-like phenotype, and boosts their stress resilience and increases the control of intracellular bacteria, as depicted in Fig.5.

**Figure 5:**
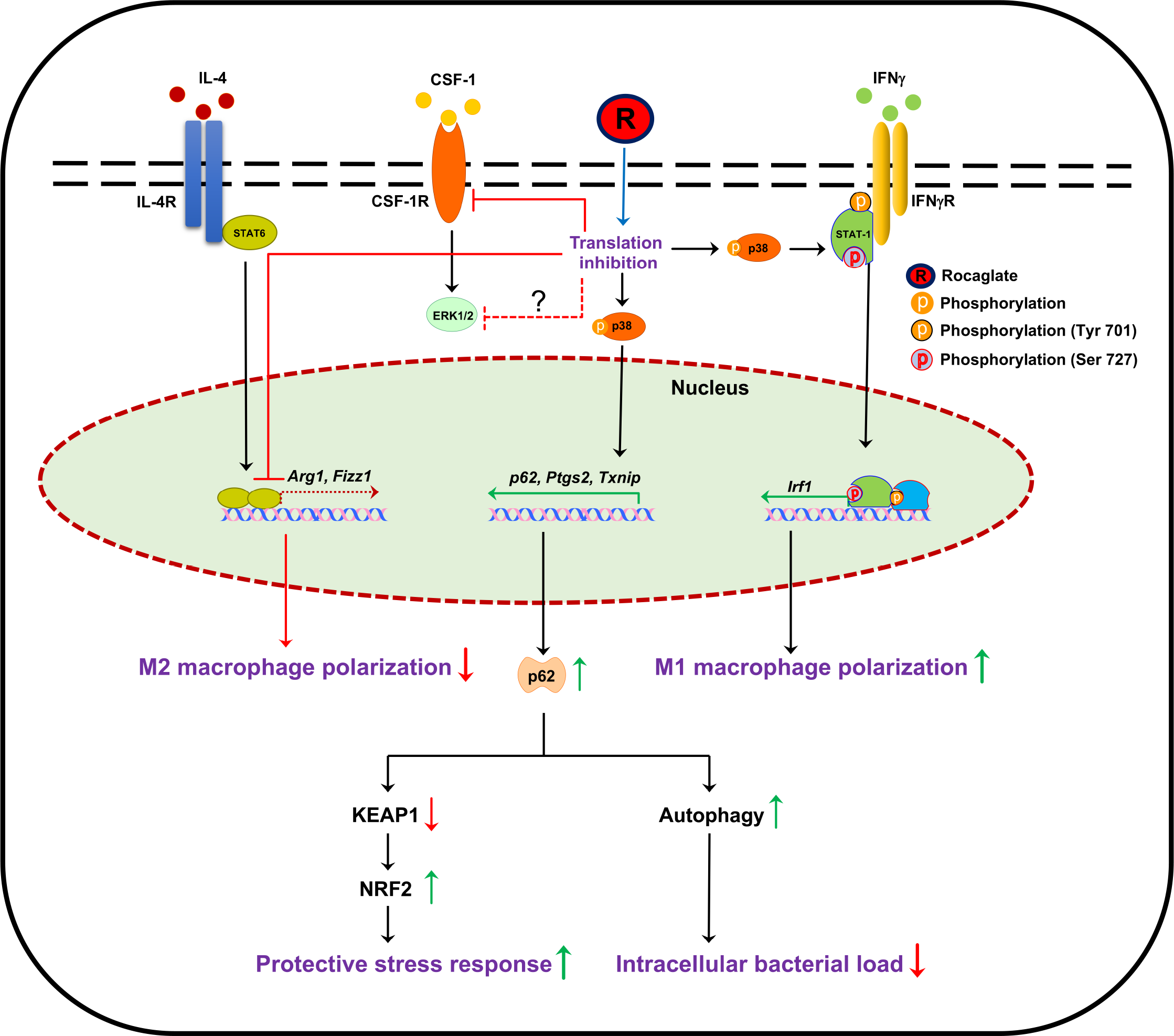
Pleiotropic effects of rocaglates’ on primary macrophages partially mimic IFNγ. Selective translation inhibition by an active rocaglate (R) induces p38-dependent and independent effects. The p38-depndent effects include synergy with low concentrations of IFNγ (via STAT1 phosphorylation at Ser 727 and upregulation of IRF1), as well as an IFNγ-independent induction of several inflammatory and stress response genes including SQSTM1/p62 protein. The p62 protein induces stress adaptation responses including anti-oxidant defense and autophagy. The p38-independent efffects include inhibition of anabolic pathways mediated by the CSF1 receptor, the IL-4 and STAT6 axis and the ERK1/2 MAPK. In toto, macrophage response to selective protein translation inhibition amounts to adaptive proteome remodelling, inhibition of cell growth and increased expression of stress response proteins. This adaptive response improves macrophage stress resilience and resistance to intracellular mycobacteria.

It is also noteworthy that biological outcomes of rocaglates on primary macrophages are distinct from their better studied effects on tumor cells. Our findings highlight that translational inhibition is necessary but not sufficient to potentiate the rocaglate effect in macrophages, since the general translational inhibitor cycloheximide and mTOR inhibitors failed to recapitulate the rocaglate-induced phenotypes. In primary macrophages, rocaglate treatment did not block translation completely. Rather, partial shutdown of translation led to reprogramming the macrophage proteome, such that proteins involved in M1-like macrophage activation and stress resilience were upregulated.

Mechanistically, synthetic and natural rocaglamide derivatives have been shown to inhibit translation by binding to a component of the eIF4F translation initiation complex – the RNA helicase eIF4A – that prevents its incorporation into the eIF4F complex (51). However, not all protein translation is equally sensitive to the inhibition of eIF4A helicase activity. Translation of mRNA species containing structural barriers, such as stem-loop structures or protein – polypurine RNA complexes, within their mRNA 5’ UTR is most sensitive to eIF4A inhibition(52, 53). Among the most eIF4A-dependent and rocaglate-sensitive transcripts in acute lymphoblastic leukemia cells were oncogene-encoding transcripts and superenhancer-associated transcription factors, such as Myc (42). Because various stressors converge on a common pathway known as Integrated Stress Response (ISR) leading to inhibition of cap-dependent protein translation, we hypothesize that the eIF4A-mediated selective downregulation of protein translation enables rapid adjustment of the anabolic pathways at translational level to match environmental inputs. Therefore, small molecules that downregulate the eIF4A RNA helicase activity may mimic those stress signals causing adaptive and reversible proteome remodeling.

We found that rocaglates potently inhibit ERK phosphorylation, a key activation step in the growth factor signal transduction relay. Phosphorylated ERK promotes cell proliferation and provides anabolic and pro-survival signals primarily via phosphorylation of transcription factors Myc and CREB. Because Myc is known to be critically important for the induction and maintenance of the “trophic” M2-like macrophage state (40, 41), ERK inhibition is likely to downregulate the Myc-dependent pathways involved in M2 polarization. According to the literature, ERK can also directly phosphorylate STAT6 and increase its DNA binding and transcriptional activities (54). We also observed that rocaglates inhibited IL-4-induced STAT6 DNA binding. Thus, profound inhibition of ERK phosphorylation by rocaglates may explain or significantly contribute to inhibition of both the STAT6- and Myc-dependent pathways of the M2 polarization and, thus, suppress both the induction and maintenance of the M2-like macrophage phenotype.

The canonical eIF4A-dependent translation inhibition by rocaglates may provide a stress-like “conservation of resources” signal that leads to the inhibition of ERK phosphorylation and, thus, to the coordinated downregulation of several anabolic pathways. Yet another mechanism may be associated with a recently described alternative target of rocaglates – prohibitin. Rocaglates have been shown to bind prohibitins directly and prevent their interaction with c-Raf, thereby, inhibiting Raf activation and Raf-MEK-ERK signaling in Jurkat tumor cell line. (22). This pathway may lead to translation inhibition by rocaglates via an eIF4A-independent, but ERK-dependent, pathway, since MEK-ERK signaling has been shown to promote cap-dependent translation via phosphorylation of eIF4E, the key translation initiation factor(22). Whether the rocaglate binding to prohibitin mediates ERK inhibition and/or translation inhibition in macrophages remains to be established(55). Our data suggest an alternative pathway of ERK down regulation in macrophages can occur via downregulation of CSF1R, a receptor for a major macrophage growth factor.

We demonstrate that rocaglates do not independently initiate the main IFNγ receptor-mediated pathway via STAT1 activation. Instead, they boost effects of IFNγ at concentrations as low as 0.05 −0.2 U/ml. Perhaps, low doses of IFNγ are necessary to provide the canonical Q701 STAT1 phosphorylation by JAK kinases at the IFNγ receptor, which is further amplified by rocaglates. This amplification of IFNγ signaling cannot be explained by reciprocal inhibition of the IL-4 - STAT6 axis by rocalates, because a specific STAT6 inhibitor AS1517499 does not recapitulate this effect (**unpublished data**). Rather, rocaglates induce a synergistic signal via p38 MAPK activation leading to additional STAT1 phosphorylation at Ser 727 (**Fig.2L**). This synergy mediates the upregulation of IRF1 mRNA expression. Obviously, the IRF1 protein translation is unimpeded by the rocaglates and, therefore, this central transcription factor in IFNγ network is significantly upregulated. Taken together, our data demonstrate that biological effects of rocaglates on primary macrophage activation are mechanistically linked to translation inhibition and mediated via modulation of MAPK signaling cascades – induction of pro-inflammatory and stress responses by p38 and suppression of anabolic pathways mediated by ERK that modulate IFNγ and IL-4 cytokine signaling via STAT1 and STAT6 modifications, respectively.

The biological effects of the rocaglate treatment are not limited to IRF1 induction. They significantly overlap with pleiotropic effects of IFNγ on macrophages, including translation inhibition, induction of autophagy and anti-oxidant defenses, as well as suppression of anabolic pathways mediated by IL-4, CSF1R and ERK (**Suppl.Table 3).** This coordinated macrophage reprogramming resembles stress preconditioning. The exposure to low, subthreshold, doses of stress prior to encountering more potent stressors was shown to increase stress resilience and prevent subsequent damage by higher doses of stressors in many biological systems. This phenomenon has been termed “hormesis” (56). In this conceptual framework, the effect of rocaglates may be viewed as anticipatory cross-protection against various stressors, including intracellular bacteria.

The dissection of signal transduction pathways activated by rocaglates demonstrate that most of their biological effects are achieved via regulatory pathways distinct from those activated by standard concentrations of IFNγ and can be interferon-independent. Therefore, rocaglates may bypass typical signal transduction initiated by the IFNγ binding to the IFNγ receptor. This property of rocaglates may be particularly important for directing macrophage polarization when the receptor signaling is disrupted by mutations, microbial factors or negative biological regulators.

The above data suggest that rocaglates may represent a novel class of HDTs with broad applications in vivo. Rocaglates are active against protozoan infections, such as malaria(57) and visceral leishmaniasis (58). In mycobacterial infections, these small molecules may penerate mycobacterial granulomas better than IFNγ and compensate for its decreased bioavailability, especially in HIV-TB co-infections. Helminth co-infections inducing the M2 macrophage polarization, increase susceptibility to mycobacterial infections (6). In these settings, the ability of rocaglates to guide monocyte/macrophages towards M1-like phenotype and inhibit triggers of the alternative, M2-like, activation may be particularly useful. Silvestrol has been shown to display a broad spectrum of antiviral activities including Corona-, Ebola-, Zika-, Picorna-, Hepatis E and Chikungunya viruses (20, 59). Recent analysis of a SARS-CoV-2 protein interaction map predicted that the rocaglate target eIF4A could be the antiviral drug target(60). Our data suggest that rocaglates may also be useful for boosting macrophage-mediated immunity and preventing bacterial pneumonias following viral infections.

Recently, it has been suggested that COVID-19 pandemics may lead to reactivation of latent TB infection and an upsurge of TB in endemic areas(61). In these settings, treatment of TB patients infected with COVID-19 with a rocaglate, may also reduce risks of TB progression, as compared to corticosteroids.

The rocaglates’ ability to direct M1-like macrophage polarization also warrants their evaluation in chronic non-infectious pathologies, where macrophages are exposed to complex, often conflicting environmental signals. Tumor associated macrophages (TAM) stimulate tumor cell proliferation, angiogenesis, metastasis and resistance to therapy. Those properties have been associated with M2-like differentiation of monocytes guided by cytokines, CSF-1 among them, within tumor microenvironments(62–64). Thus, in solid tumors, rocaglates may target both the tumor cells decreasing their proliferation and myeloid cells that comprise the immune and inflammatory milieu. In asthma, allergic reactions may be associated with chronic bacterial infections. Currently, asthma therapy relies on immune suppression by corticosteroids. However, prolonged treatment with steroids has numerous undesirable side effects, including chronic pulmonary infections caused by opportunistic mycobacteria and fungi that further exacerbate pulmonary pathology (65). The important theoretical advantage of rocaglates as compared to corticosteroids is that rocaglates would shift the balance of macrophage activation towards M1-like phenotype without suppressing mechanisms of macrophage resistance to bacterial and viral infections. Thus, rocaglates may represent a novel class of HDTs broadly applicable to diseases associated with hypofunction of type 1 and/or hyperactivation of type 2 immunity, e.g., chronic bacterial infections, allergies and, possibly, certain tumors.

## Materials and Methods

### Mice

C57BL/6 J mice were obtained from the Jackson Laboratory (USA). All experiments were performed with the full knowledge and approval of the Standing Committee on Animals at Boston University (IACUC protocol number AN15276).

### Bacterial strains

The wild type *M. tuberculosis* H37Rv, *M. bovis* BCG Pasteur and recombinant *M. bovis* BCG expressing *gfp* strains were grown at 37°C in Middlebrook 7H9 broth (BD Biosciences) or on 7H10 agar plates (BD Biosciences), respectively. The both solid and liquid media contained contained glycerol (0.5% v/v) and Tween 80 (0.05%). The MB7H9 broth was enriched using 10% ADC and MB7H10 agar was enriched using with 10% OADC. The *M. bovis* BCG strain expressing *gfp* were grown with 50 μg/mL Hygromycin B.

### BMDM culture

Isolation of mouse bone marrow and culture of BMDMs were carried out as previously described(66).

### Gel shift assay

The nuclear extracts were prepared by nuclear extraction kits (Signosis Inc). Gel shift assays were done with EMSA kits (Signosis Inc). 5 μg nuclear extracts were incubated with 1× binding buffer and biotin-labeled probe for 30 min at room temperature. The samples were electrophoresed on a 6 % polyacrylamide gel in 0.5 % TBE at 120 V for 45 min and transferred onto a nylon membrane in 0.5 % TBE at 300 mA for 1 h. After transfer and UV cross-linking, the membrane was detected with Streptavidin–HRP.

### Immunoblotting

Equal amounts of protein extracts were separated by SDS-PAGE and transferred to PVDF membrane (Millipore). Bands were detected with enhanced chemiluminescence (ECL) kit (Perkin Elmer).

### RNA isolation and quantitative PCR

Total RNA was isolated using the RNeasy Plus mini kit (Qiagen). cDNA synthesis was performed using the SuperScript II (Invitrogen). Quantitative real-time RT-PCR (qRT-PCR) was performed with the GoTaq qPCR Mastermix (Promega) using the CFX-90 real-time PCR System (Bio-Rad).

### Hoechst/PI staining method for cell cytotoxicity

For cell viability assays BMDMs were plated in 96 well tissue culture plates. The supernatant was aspirated and to each well 100□μl PBS containing Hoechst (Invitrogen, 10□μM) and PI (Calbiochem, 2□μM) were added. The % of total and dead cells was calculated by automated cell cytometer.

### Reporter assay for measuring translation inhibition

293TR-Fluc cells, a gift of Dr. Whitesell (16) were grown to confluence. Compounds were added at different dilutions and kept for 18□h. 100□ul of Nanolight Firefly Luc Assay reagent was added to the wells and the luminescence was measured using a Tecan-plate reader after 2□mins. Cells treated with DMSO served as negative controls and with 100 μg/mL cycloheximide served as positive controls. Percentage of translation inhibition for each compound was calculated in triplicates from two independent experiments.

### Puromycin incorporation assay

Puromycin labelling for measuring the intensity of translation was performed as described(67). In brief, 5 μg/ml puromycin (Sigma,) was added in the culture medium and incubated for 1 h at 37 °C and 5% CO_2_. Immunoblotting was performed with the anti-puromycin antibody clone12D10 (1:10000) (Millipore).

### Phagosome–lysosome fusion assay

The cells were pretreated with CMLD010536 (25 nM) for 6 h and subsequently infected with *M. bovis* BCG expressing *gfp* (BCG:*gfp*) for 1 h at MOI 1. To remove the extracellular bacteria, the cells were treated with 200 ug/ml of amikacin for 1 h. After 1 h, 24 h and 48 h post-infection, cells were stained with 200 nM of LysoTracker red dye for 1 h at 37 °C and analyzed using Zeiss LSM 710-Live Duo scan confocal microscope. The images were processed using FIJI software and the percentage of co-localization of BCG:*gfp* containing phagosomes with LysoTracker red dye was calculated by dividing the number of co-localized phagosomes by the total number of phagosomes. At least 50 cells are measured at each condition and the mean ± SEM (*p value* was ??<??>0.05) was calculated using GraphPad Prism.

### Macrophage infection and determination of intracellular bacterial loads

BMDM were infected with the mycobacteria in 96 well plate at preferred multiplicity of infection (MOI) by suspending bacteria in BMDM specific medium. The infected cells were centrifuged for 5 min at 200xg and incubated at 37°C with 5% CO_2_ for 1 h and extracellular bacteria were killed off by incubating the cells in medium containing 200 μg/ml amikacin for 1 h. Further BMDM were washed 3 times with 1% PBS containing 2% FBS and incubated at 37°C with 5% CO2 until further analysis. The intracellular bacterial load was determined using quantitative real time PCR and standard CFU methods. BMDM were lysed using 0.05% Tween 80 in 1X PBS for 5 min at room temperature. The lysates were serially diluted and used for plating on Middlebrook 7H10 agar. DNA for quantitative PCR method was isolated using modified Mtb lysis buffer and magnetic bead purification, as described in detail in Yabaji *et al*., submitted for publication).

### Statistical analysis

The densitometric analyses were performed using ImageJ 2.0 software and all the graphs were plotted using Graphpad Prism 8. For multiple treatment experiments, Students’s unpaired t-test and one-way ANOVA analysis was performed in Graphpad Prism 8. The *P values* of ≤0.05 were considered statistically significant.

## Acknowledgments

This work was supported by the National Institutes of Health, award numbers R33 AI105944-04 and R01 HL133190-01 (IK), 1R01GM120272 (ARI) and R01CA218500 (ARI). The authors are grateful to Dr. John Connor for helpful discussions.

The authors declare no competing interests.

## References

1. Wallis RS, and Hafner R. Advancing host-directed therapy for tuberculosis. Nat Rev Immunol. 2015;15(4):255–63.

2. Heitmann L, Abad Dar M, Schreiber T, Erdmann H, Behrends J, McKenzie AN, et al. The IL-13/IL-4Ralpha axis is involved in tuberculosis-associated pathology. J Pathol. 2014;234(3):338–50.

3. Chanmee T, Ontong P, Konno K, and Itano N. Tumor-associated macrophages as major players in the tumor microenvironment. Cancers (Basel). 2014;6(3):1670–90.

4. Sica A, and Mantovani A. Macrophage plasticity and polarization: in vivo veritas. J Clin Invest. 2012;122(3):787–95.

5. Potian JA, Rafi W, Bhatt K, McBride A, Gause WC, and Salgame P. Preexisting helminth infection induces inhibition of innate pulmonary anti-tuberculosis defense by engaging the IL-4 receptor pathway. J Exp Med. 2011;208(9):1863–74.

6. Salgame P, Yap GS, and Gause WC. Effect of helminth-induced immunity on infections with microbial pathogens. Nat Immunol. 2013;14(11):1118–26.

7. Harris J, De Haro SA, Master SS, Keane J, Roberts EA, Delgado M, et al. T helper 2 cytokines inhibit autophagic control of intracellular Mycobacterium tuberculosis. Immunity. 2007;27(3):505–17.

8. Patel SY, Ding L, Brown MR, Lantz L, Gay T, Cohen S, et al. Anti-IFN-gamma autoantibodies in disseminated nontuberculous mycobacterial infections. J Immunol. 2005;175(7):4769–76.

9. Browne SK, and Holland SM. Anticytokine autoantibodies in infectious diseases: pathogenesis and mechanisms. Lancet Infect Dis. 2010;10(12):875–85.

10. Venkataraman C, Leung S, Salvekar A, Mano H, and Schindler U. Repression of IL-4-induced gene expression by IFN-gamma requires Stat1 activation. J Immunol. 1999;162(7):4053–61.

11. Cadena AM, Fortune SM, and Flynn JL. Heterogeneity in tuberculosis. Nat Rev Immunol. 2017;17(11):691–702.

12. Carow B, Hauling T, Qian X, Kramnik I, Nilsson M, and Rottenberg ME. Spatial and temporal localization of immune transcripts defines hallmarks and diversity in the tuberculosis granuloma. Nat Commun. 2019;10(1):1823.

13. Marino S, Cilfone NA, Mattila JT, Linderman JJ, Flynn JL, and Kirschner DE. Macrophage polarization drives granuloma outcome during Mycobacterium tuberculosis infection. Infect Immun. 2015;83(1):324–38.

14. Sakai S, Mayer-Barber KD, and Barber DL. Defining features of protective CD4 T cell responses to Mycobacterium tuberculosis. Curr Opin Immunol. 2014;29:137–42.

15. Bhattacharya B, Chatterjee S, Devine WG, Kobzik L, Beeler AB, Porco JA, Jr., et al. Fine-tuning of macrophage activation using synthetic rocaglate derivatives. Sci Rep. 2016;6:24409.

16. Santagata S, Mendillo ML, Tang YC, Subramanian A, Perley CC, Roche SP, et al. Tight coordination of protein translation and HSF1 activation supports the anabolic malignant state. Science. 2013;341(6143):1238303.

17. Hwang BY, Su BN, Chai H, Mi Q, Kardono LB, Afriastini JJ, et al. Silvestrol and episilvestrol, potential anticancer rocaglate derivatives from Aglaia silvestris. J Org Chem. 2004;69(10):3350–8.

18. Kim S, Hwang BY, Su BN, Chai H, Mi Q, Kinghorn AD, et al. Silvestrol, a potential anticancer rocaglate derivative from Aglaia foveolata, induces apoptosis in LNCaP cells through the mitochondrial/apoptosome pathway without activation of executioner caspase-3 or −7. Anticancer Res. 2007;27(4B):2175–83.

19. Kim S, Salim AA, Swanson SM, and Kinghorn AD. Potential of cyclopenta[b]benzofurans from Aglaia species in cancer chemotherapy. Anticancer Agents Med Chem. 2006;6(4):319–45.

20. Müller C, Schulte FW, Lange-Grünweller K, Obermann W, Madhugiri R, Pleschka S, et al. Broad-spectrum antiviral activity of the eIF4A inhibitor silvestrol against corona- and picornaviruses. Antiviral research. 2018;150:123–9.

21. Bordeleau ME, Robert F, Gerard B, Lindqvist L, Chen SM, Wendel HG, et al. Therapeutic suppression of translation initiation modulates chemosensitivity in a mouse lymphoma model. J Clin Invest. 2008;118(7):2651–60.

22. Polier G, Neumann J, Thuaud F, Ribeiro N, Gelhaus C, Schmidt H, et al. The natural anticancer compounds rocaglamides inhibit the Raf-MEK-ERK pathway by targeting prohibitin 1 and 2. Chem Biol. 2012;19(9):1093–104.

23. Rodrigo CM, Cencic R, Roche SP, Pelletier J, and Porco Jr. JA. Synthesis of Rocaglamide Hydroxamates and Related Compounds as Eukaryotic Translation Inhibitors: Synthetic and Biological Studies. Journal of Medicinal Chemistry. 2012;55(1):558–62.

24. Katsuragi Y, Ichimura Y, and Komatsu M. p62/SQSTM1 functions as a signaling hub and an autophagy adaptor. FEBS J. 2015;282(24):4672–8.

25. Moscat J, and Diaz-Meco MT. Feedback on fat: p62-mTORC1-autophagy connections. Cell. 2011;147(4):724–7.

26. Knaevelsrud H, and Simonsen A. Fighting disease by selective autophagy of aggregate-prone proteins. FEBS Lett. 2010;584(12):2635–45.

27. Johansen T, and Lamark T. Selective autophagy mediated by autophagic adapter proteins. Autophagy. 2011;7(3):279–96.

28. Taguchi K, Motohashi H, and Yamamoto M. Molecular mechanisms of the Keap1-Nrf2 pathway in stress response and cancer evolution. Genes Cells. 2011;16(2):123–40.

29. Jaramillo MC, and Zhang DD. The emerging role of the Nrf2-Keap1 signaling pathway in cancer. Genes Dev. 2013;27(20):2179–91.

30. Padmanabhan B, Tong KI, Ohta T, Nakamura Y, Scharlock M, Ohtsuji M, et al. Structural basis for defects of Keap1 activity provoked by its point mutations in lung cancer. Mol Cell. 2006;21(5):689–700.

31. Fukutomi T, Takagi K, Mizushima T, Ohuchi N, and Yamamoto M. Kinetic, thermodynamic, and structural characterizations of the association between Nrf2-DLGex degron and Keap1. Mol Cell Biol. 2014;34(5):832–46.

32. Komatsu M, Kurokawa H, Waguri S, Taguchi K, Kobayashi A, Ichimura Y, et al. The selective autophagy substrate p62 activates the stress responsive transcription factor Nrf2 through inactivation of Keap1. Nat Cell Biol. 2010;12(3):213–23.

33. Lau A, Wang XJ, Zhao F, Villeneuve NF, Wu T, Jiang T, et al. A noncanonical mechanism of Nrf2 activation by autophagy deficiency: direct interaction between Keap1 and p62. Mol Cell Biol. 2010;30(13):3275–85.

34. Meraz MA, White JM, Sheehan KC, Bach EA, Rodig SJ, Dighe AS, et al. Targeted disruption of the Stat1 gene in mice reveals unexpected physiologic specificity in the JAK-STAT signaling pathway. Cell. 1996;84(3):431–42.

35. Decker T, Kovarik P, and Meinke A. GAS elements: a few nucleotides with a major impact on cytokine-induced gene expression. J Interferon Cytokine Res. 1997;17(3):121–34.

36. Dudley AC, Thomas D, Best J, and Jenkins A. The STATs in cell stress-type responses. Cell Commun Signal. 2004;2(1):8.

37. Wen Z, Zhong Z, and Darnell JE, Jr. Maximal activation of transcription by Stat1 and Stat3 requires both tyrosine and serine phosphorylation. Cell. 1995;82(2):241–50.

38. Kovarik P, Stoiber D, Novy M, and Decker T. Stat1 combines signals derived from IFN-gamma and LPS receptors during macrophage activation. EMBO J. 1998;17(13):3660–8.

39. Goh KC, Haque SJ, and Williams BR. p38 MAP kinase is required for STAT1 serine phosphorylation and transcriptional activation induced by interferons. EMBO J. 1999;18(20):5601–8.

40. Pello OM. Macrophages and c-Myc cross paths. Oncoimmunology. 2016;5(6):e1151991.

41. Pello OM, De Pizzol M, Mirolo M, Soucek L, Zammataro L, Amabile A, et al. Role of c-MYC in alternative activation of human macrophages and tumor-associated macrophage biology. Blood. 2012;119(2):411–21.

42. Wolfe AL, Singh K, Zhong Y, Drewe P, Rajasekhar VK, Sanghvi VR, et al. RNA G-quadruplexes cause eIF4A-dependent oncogene translation in cancer. Nature. 2014;513(7516):65–70.

43. Rogov V, Dotsch V, Johansen T, and Kirkin V. Interactions between autophagy receptors and ubiquitin-like proteins form the molecular basis for selective autophagy. Mol Cell. 2014;53(2):167–78.

44. Zheng YT, Shahnazari S, Brech A, Lamark T, Johansen T, and Brumell JH. The adaptor protein p62/SQSTM1 targets invading bacteria to the autophagy pathway. J Immunol. 2009;183(9):5909–16.

45. Chitu V, Yeung YG, Yu W, Nandi S, and Stanley ER. Measurement of Macrophage Growth and Differentiation. Current protocols in immunology. 2011;92(1):14.20.1–14.20.6.

46. Vergadi E, Ieronymaki E, Lyroni K, Vaporidi K, and Tsatsanis C. Akt Signaling Pathway in Macrophage Activation and M1/M2 Polarization. The Journal of Immunology. 2017;198(3):1006–14.

47. Mattila JT, Ojo OO, Kepka-Lenhart D, Marino S, Kim JH, Eum SY, et al. Microenvironments in Tuberculous Granulomas Are Delineated by Distinct Populations of Macrophage Subsets and Expression of Nitric Oxide Synthase and Arginase Isoforms. Journal of immunology (Baltimore, Md: 1950). 2013;191(2):773–84.

48. Yang Z, Huang Y-CT, Koziel H, de Crom R, Ruetten H, Wohlfart P, et al. Female resistance to pneumonia identifies lung macrophage nitric oxide synthase-3 as a therapeutic target. eLife. 2014;3:265.

49. Dockrell DH, Marriott HM, Prince LR, Ridger VC, Ince PG, Hellewell PG, et al. Alveolar macrophage apoptosis contributes to pneumococcal clearance in a resolving model of pulmonary infection. J Immunol. 2003;171(10):5380–8.

50. Gordon SB, Irving GR, Lawson RA, Lee ME, and Read RC. Intracellular trafficking and killing of Streptococcus pneumoniae by human alveolar macrophages are influenced by opsonins. Infect Immun. 2000;68(4):2286–93.

51. Cencic R, Carrier M, Galicia-Vazquez G, Bordeleau ME, Sukarieh R, Bourdeau A, et al. Antitumor activity and mechanism of action of the cyclopenta[b]benzofuran, silvestrol. PLoS One. 2009;4(4):e5223.

52. Iwasaki S, Floor SN, and Ingolia NT. Rocaglates convert DEAD-box protein eIF4A into a sequence-selective translational repressor. Nature. 2016;534(7608):558–61.

53. Iwasaki S, Iwasaki W, Takahashi M, Sakamoto A, Watanabe C, Shichino Y, et al. The Translation Inhibitor Rocaglamide Targets a Bimolecular Cavity between eIF4A and Polypurine RNA. Molecular cell. 2019;73(4):738–48.e9.

54. So EY, Oh J, Jang JY, Kim JH, and Lee CE. Ras/Erk pathway positively regulates Jak1/STAT6 activity and IL-4 gene expression in Jurkat T cells. Mol Immunol. 2007;44(13):3416–26.

55. Chu J, Galicia-Vázquez G, Cencic R, Mills JR, Katigbak A, Porco JA, et al. CRISPR-Mediated Drug-Target Validation Reveals Selective Pharmacological Inhibition of the RNA Helicase, eIF4A. CELREP. 2016;15(11):2340–7.

56. Calabrese EJ. Hormesis: why it is important to toxicology and toxicologists. Environ Toxicol Chem. 2008;27(7):1451–74.

57. Langlais D, Cencic R, Moradin N, Kennedy JM, Ayi K, Brown LE, et al. Rocaglates as dual-targeting agents for experimental cerebral malaria. Proceedings of the National Academy of Sciences of the United States of America. 2018;8:201713000.

58. Chaparro V, Leroux L-P, Masvidal L, Lorent J, Graber TE, Zimmermann A, et al. Translational profiling of macrophages infected with Leishmania donovani identifies mTOR- and eIF4A-sensitive immune-related transcripts. PLoS pathogens. 2020;16(6):e1008291.

59. Blum L, Geisslinger G, Parnham MJ, Grünweller A, and Schiffmann S. Natural antiviral compound silvestrol modulates human monocyte derived macrophages and dendritic cells. Journal of Cellular and Molecular Medicine. 2020;24(12):6988–99.

60. Gordon DE, Jang GM, Bouhaddou M, Xu J, Obernier K, White KM, et al. A SARS-CoV-2 protein interaction map reveals targets for drug repurposing. Nature. 2020:1–13.

61. Nordling L. HIV and TB increase death risk from COVID-19, study finds—but not by much. Science (New York, NY). 2020.

62. Schmid MC, and Varner JA. Myeloid cells in the tumor microenvironment: modulation of tumor angiogenesis and tumor inflammation. J Oncol. 2010;2010:201026.

63. Murray PJ. Nonresolving□macrophage mediated inflammation in malignancy. FEBS Journal. 2018;285(4):641–53.

64. Grivennikov SI, Greten FR, and Karin M. Immunity, inflammation, and cancer. Cell. 2010;140(6):883–99.

65. Fritscher LG, Marras TK, Bradi AC, Fritscher CC, Balter MS, and Chapman KR. Nontuberculous Mycobacterial Infection as a Cause of Difficult-to-Control Asthma: A Case-Control Study. Chest. 2011;139(1):23–7.

66. Pan H, Mostoslavsky G, Eruslanov E, Kotton DN, and Kramnik I. Dual-promoter lentiviral system allows inducible expression of noxious proteins in macrophages. J Immunol Methods. 2008;329(1-2):31–44.

67. Schmidt EK, Clavarino G, Ceppi M, and Pierre P. SUnSET, a nonradioactive method to monitor protein synthesis. Nature methods. 2009;6(4):275–7.

