## Supplemental Figures for "Channeling macrophage polarization via selective translation inhibition by rocaglates increases macrophage resistance to Mycobacterium tuberculosis"

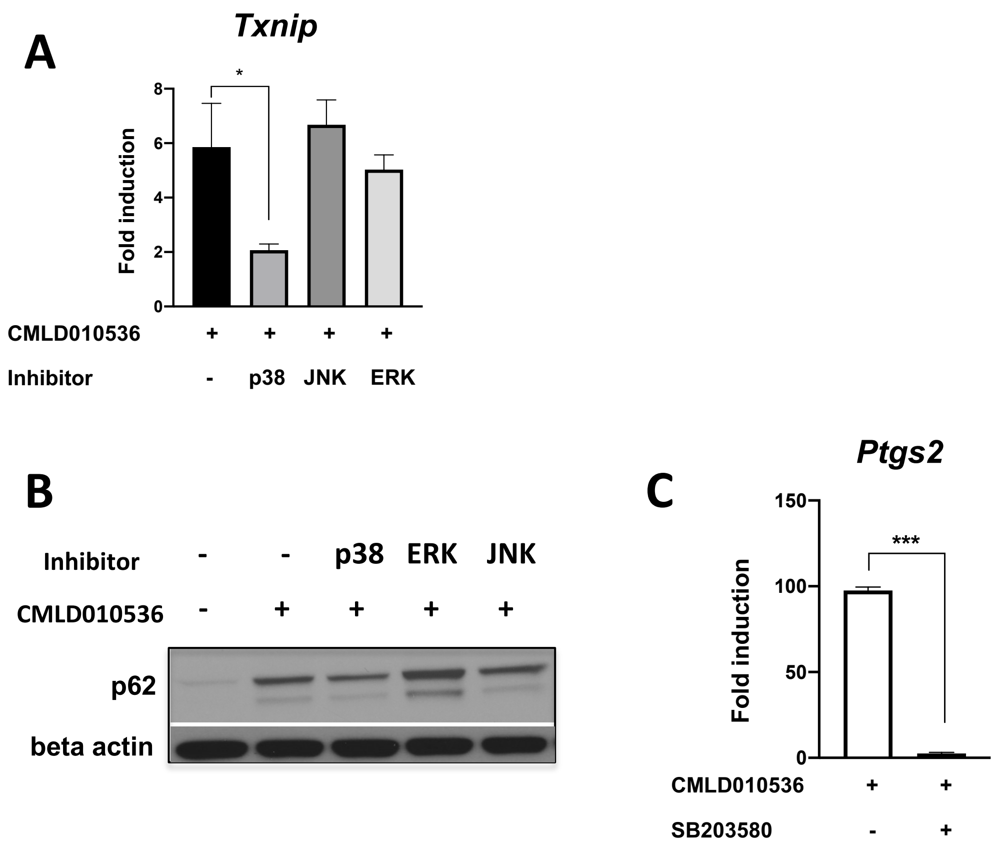


**Supplementary Figure 1.**

**(A)** Macrophages were pretreated with 10 µM MAPK kinase inhibitors (SB203580, SB600125 and U0126) for 30 min followed by treatment with 100 nM CMLD010536 in the presence of the inhibitors for additional 24 h. The mRNA expression for *Txnip* was measured by qRT-PCR.

**(B)** Macrophages were pretreated with 10 µM MAPK kinase inhibitors (SB203580, SB600125 and U0126) for 30 mins followed by treatment with 100 nM CMLD010536 in the presence of the inhibitors for additional 24 h. Expression of *p62* protein was measured by Western blot and compared to cells treated with CMLD010536 alone.

**(C)** Macrophages were pretreated with 10 mM p38 inhibitor (SB203580) for 30 mins followed by treatment with 0.3 mM CMLD010536 for additional 4 h. Expression of *Ptgs2* mRNA was measured by qRT-PCR. The data are represented as mean ± SEM and *p value* ≤ 0.05 was considered statistically significant.

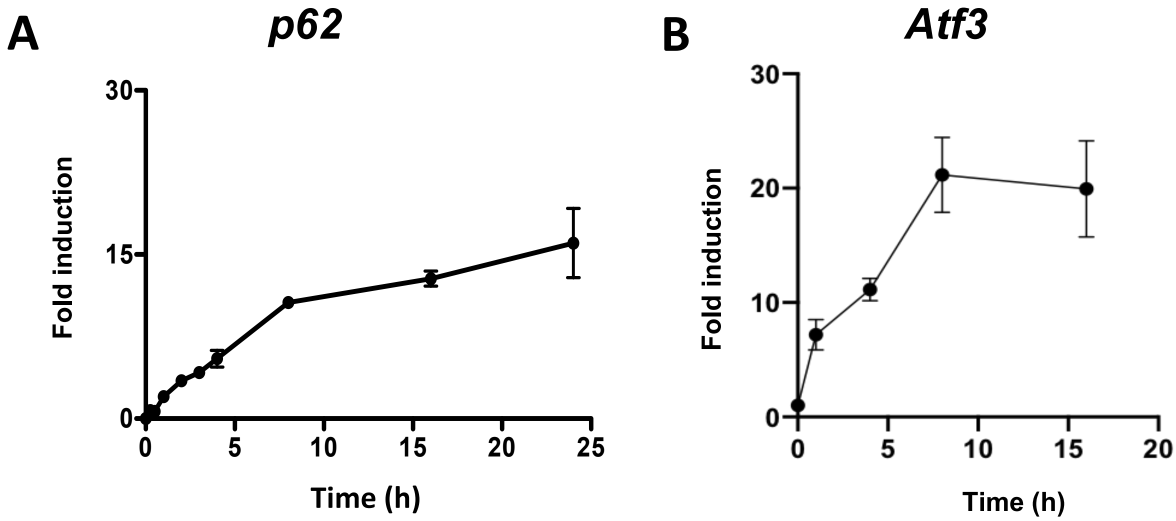


**Supplementary Figure 2.**

**(A) The kinetics of p62 induction by the active rocaglate.** BMDMs were treated with 100 nM CMLD010536 for the indicated period of time and induction of *p62* expression was measured by qRT-PCR.

(**B**) BMDMs were treated with 100 nM CMLD010536 for the indicated time and the *Atf3* expression was measured by qRT-PCR. The data are represented as mean ± SEM and *p value* ≤ 0.05 was considered statistically significant.

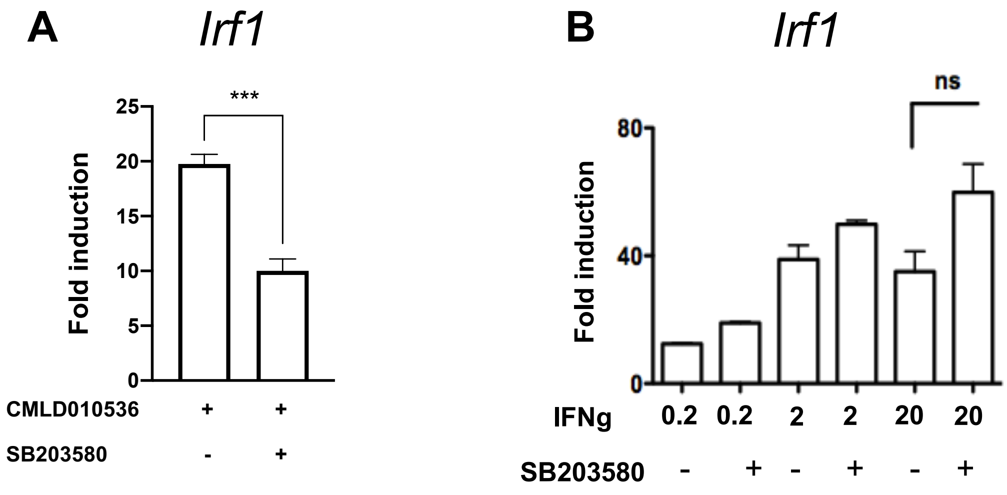


**Supplementary Figure 3.**

**(A) p38 contributes to the *Irf1* co-stimulation by rocaglates.** Macrophages were pretreated with 10 μM of p38 MAPK inhibitor (SB203580) for 30 min followed by treatment with 0.3 μM CMLD010536 and 0.2 U/ml IFNγ for additional 4 h. The *Irf1* mRNA expression was measured by qRT-PCR.

**(B) p38 does not contribute to the *Irf1* induction by IFNγ.** Macrophages were pre-treated with 10 μM SB203580 for 30 min and subsequently treated with 0.2, 2 or 20 U/ml IFNγ for 4 h and *Irf1* expression was measured by qRT-PCR. The data are represented as mean ± SEM, *p value* ≤ 0.05 was considered statistically significant.

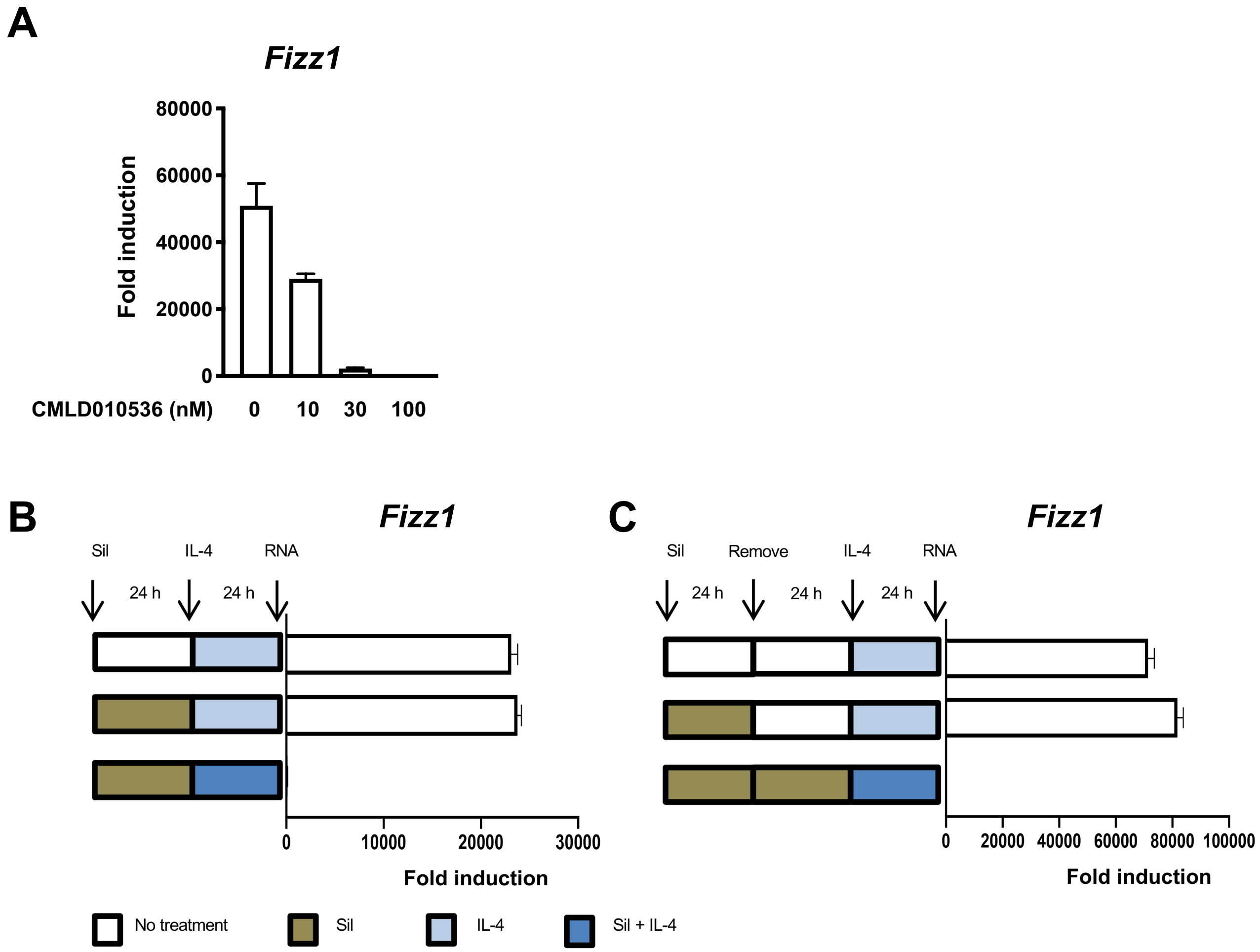


**Supplementary Figure 4:**

**(A)** BMDMs were pretreated with indicated concentrations of CMLD010536 for 4 h, followed by a treatment with 25 ng/ml IL4 for additional 20 h. Suppression of *Fizz1* mRNA was measured using qRT-PCR.

**(B)** BMDMs were pretreated with 100 nM silvestrol for 24 h followed by treatment with IL-4 alone (100 ng/ml) or IL-4 plus silvestrol (100 ng/ml and 100 nM respectively) for additional 24 h. The expression of IL-4 regulated gene, *Fizz1* was measured using qRT-PCR. The cells without pretreatment were used as control.

**(C)** BMDMs were pretreated with 100 nM silvestrol for 24 h, washed, rested for 24 h and stimulated with IL-4 (100 ng/ml) for an additional 24 h period. The *Fizz1* mRNA was measured using qRT-PCR. Cell cultured in the presence or absence of silvestrol for the duration of the experiment served as positive and negative controls.

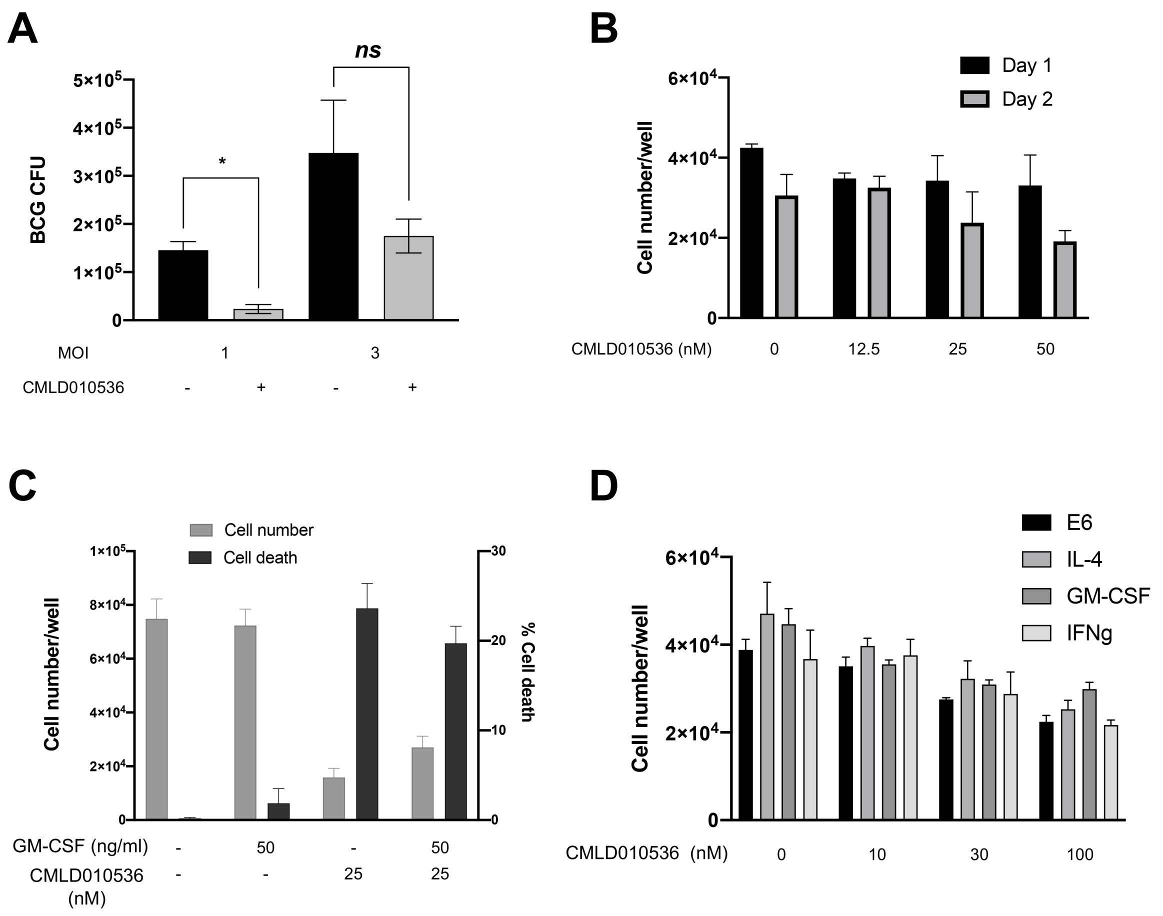


**Supplementary Figure 5.**

(**A)** BMDMs were pretreated with CMLD010536 (50 nM) for 24 h and subsequently infected with *M. bovis* BCG for 24 h at MOI 1 or 3. The intracellular BCG loads were enumerated by CFU on 7H10 agar.

**(B)** BMDMs were treated with CMLD010536 (12.5, 25 and 50 nM) for 24 and 48 h and the total cell number was determined using automated cytometry.

**(C)** BMDMs were pretreated with GM-CSF (50 ng/mL) for 16 h and subsequently infected with Mtb at MOI 1 followed by treatment with 25 nM CMLD010536 either alone or in combination with GM-CSF (50 ng/mL) for 72 hrs. The total cell number (left Y axis) and percentage cell death (right Y axis) were determined using automated cytometry (Celigo). The dotted line in graph indicates the cell number before treatement (day 0).

**(D).** BMDM were treated with different concentrations of CMLD010536 alone or in combination with IL-4 (100 ng/mL), GM-CSF (50 ng/mL) or IFNγ (100U/mL) for 24 h and the total cell numbers were determined as above. The data are presented as mean ± SEM and *p value* ≤ 0.05 was considered statistically significant.

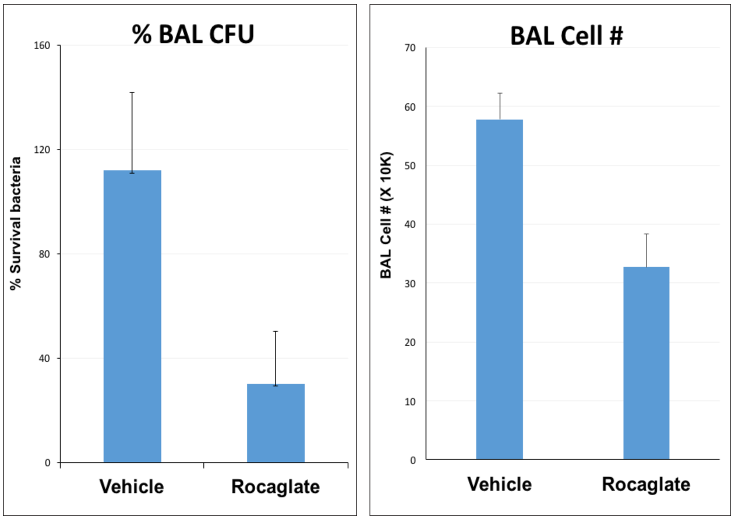


**Supplementary Figure 6**:
(A) D-1 IGS outbred male mice were injected i.p. with CMLD010536 (25 mcg/kg) or the vehicle control at 48, 24 and 0 h before respiratory infection with *Strep.pneumoniae*. At 24 h p.i., BAL fluid was harvested to determine viable bacterial counts (CFU, left panel) and total BAL cells (right panel).
