## Supplemental Table 1 for "Channeling macrophage polarization via selective translation inhibition by rocaglates increases macrophage resistance to Mycobacterium tuberculosis"

**Supplementary Table 1**

| Number | Compound ID | Structure | Translational Inhibition $IC_{50}$ (μM) | IRF1 (0.3 μM Compound) in presence of 0.1 U IFNg (24 hrs) |
|---|---|---|---|---|
| 1 | CMLD007564 | 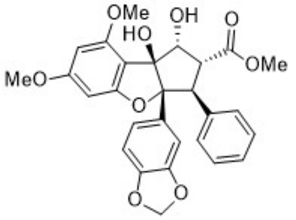 | 0.914 | 3.3 |
| 2 | CMLD007565 | 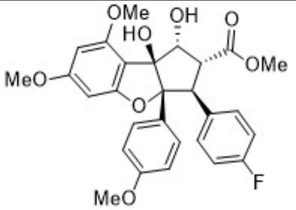 | 0.808 | 2.5 |
| 3 | CMLD010021 | 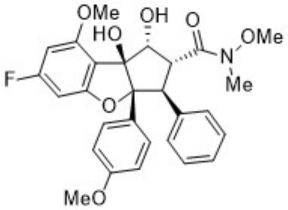 | 0.582 | 4.2 |
| 4 | CMLD010483 | 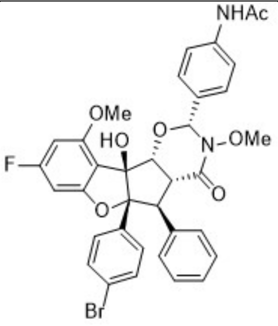 | 0.108 | 13.3 |
| 5 | CMLD010506 (Aglaroxin C) | 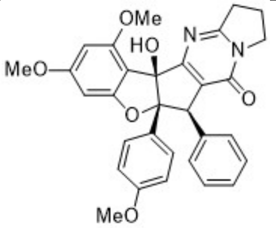 | 0.044 | 22.1 |

| Number | Compound ID | Structure | Translational Inhibition $IC_{50}$ (μM) | IRF1 (0.3 μM Compound) in presence of 0.1 U IFNg (24 hrs) |
|---|---|---|---|---|
| 6 | CMLD010510 | 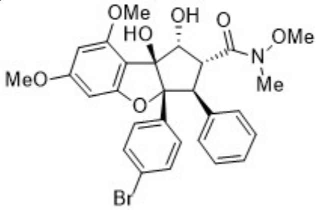 | 0.098 | 18.3 |
| 7 | CMLD010512 | 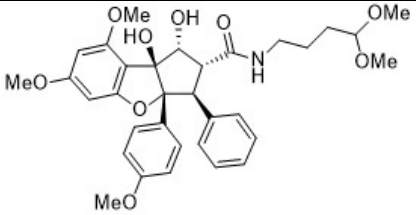 | 0.032 | 9.5 |
| 8 | CMLD010515 | 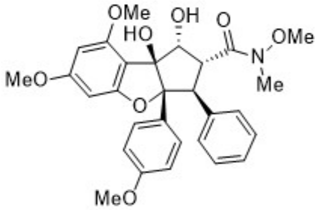 | 0.009 | 36.3 |
| 9 | CMLD010517 | 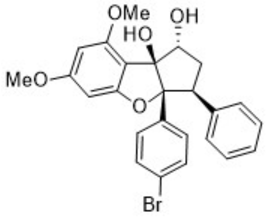 | 0.055 | 16.7 |
| 10 | CMLD010535 | 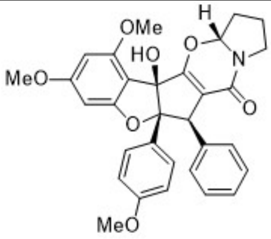 | 0.054 | 25.4 |
| 11 | CMLD010536 | 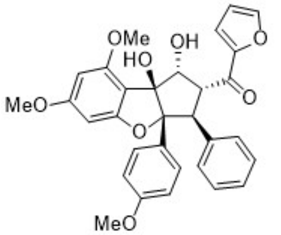 | 0.020 | 38.4 |
| 12 | CMLD010582 | 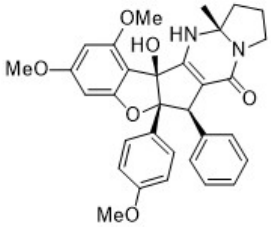 | 0.106 | 4.8 |

| Number | Compound ID | Structure | Translational Inhibition $IC_{50}$ (μM) | IRF1 (0.3 μM Compound) in presence of 0.1 U IFNg (24 hrs) |
|---|---|---|---|---|
| 13 | CMLD009433 | 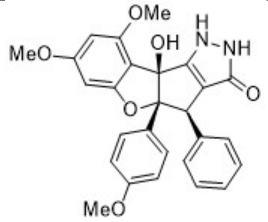 | 0.053 | 28.5 |
| 14 | silvestrol | 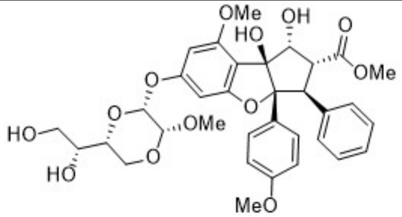 | 0.035 | 42.4 |
| 15 | CMLD011403 | 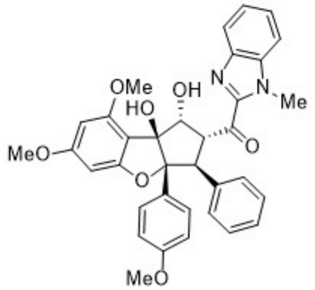 | 0.230 | 13.8 |
| 16 | CMLD011404 | 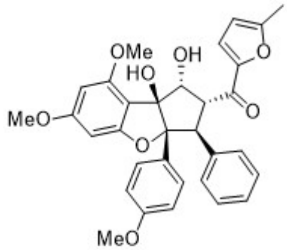 | 0.020 | 41.7 |
| 17 | CMLD011405 | 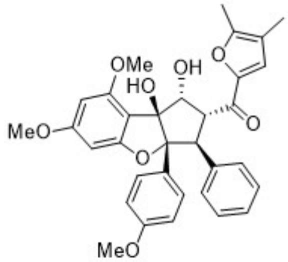 | 0.014 | 33.9 |
| 18 | CMLD011406 | 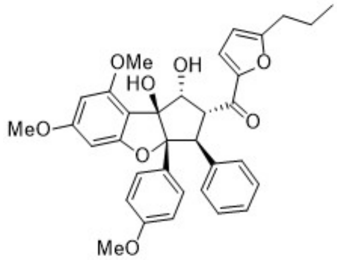 | 0.034 | 19.8 |

| Number | Compound ID | Structure | Translational Inhibition $IC_{50}$ (µM) | IRF1 (0.3 µM Compound) in presence of 0.1 U IFNg (24 hrs) |
|---|---|---|---|---|
| 19 | CMLD011407 | 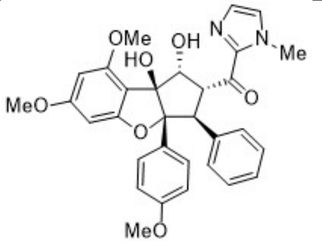 | 0.010 | 26.1 |
| 20 | CMLD012172 | 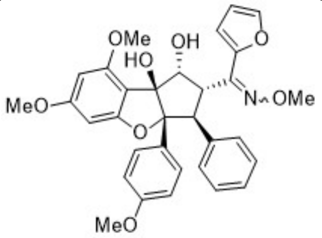 | 0.130 | 7.5 |
| 21 | CMLD011408 | 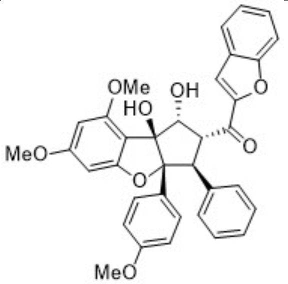 | 0.020 | 35.9 |
| 22 | CMLD011409 | 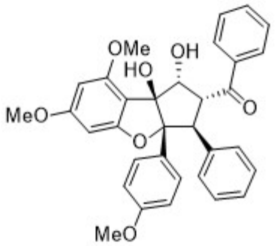 | 0.020 | 13.7 |
| 23 | CMLD011410 | 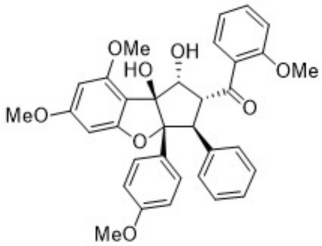 | 0.030 | 29.9 |
