## Supplemental Table 2 for "Channeling macrophage polarization via selective translation inhibition by rocaglates increases macrophage resistance to Mycobacterium tuberculosis"

### Supplementary Table 2. Key Proteins Upregulated by CMLD010536

| ID | ID2 | ACCESSION | Name | Gene |
|---|---|---|---|---|
| **P39688** | P39688 | FYN_MOUSE | Tyrosine-protein kinase Fyn (EC 2.7.10.2) (Proto-oncogene c-Fyn) (p59-Fyn) | Fyn |
| **A2AQU8** | A2AQU8 | A2AQU8_MOUSE | Sulfiredoxin-1 | Srxn1 |
| **Q60765** | Q60765 | ATF3_MOUSE | Cyclic AMP-dependent transcription factor ATF-3 (cAMP-dependent transcription factor ATF-3) (Activating transcription factor 3) (Transcription factor LRG-21) | Atf3 |
| **Q64337** | Q64337 | SQSTM_MOUSE | Sequestosome-1 (STONE14) (Ubiquitin-binding protein p62) | Sqstm1 |
| **Q99MK9-2** | Q99MK9-2 | RASF1_MOUSE | Ras association domain-containing protein 1 (Protein 123F2) | |
| **A6H663** | A6H663 | A6H663_MOUSE | BCL2-associated athanogene 3 | Bag3 |
| **Q6GTM0** | Q6GTM0 | Q6GTM0_MOUSE | Ifit2 protein (Interferon-induced protein with tetratricopeptide repeats 2) | Ifit2 |
| **Q6GT24** | Q6GT24 | Q6GT24_MOUSE | Peroxiredoxin 6 (Peroxiredoxin-6) | Prdx6 |
| **Q4FJY6** | Q4FJY6 | Q4FJY6_MOUSE | Ifit3 protein | Ifit3 |
| **P10630-2** | P10630-2 | IF4A2_MOUSE | Eukaryotic initiation factor 4A-II (eIF-4A-II) (eIF4A-II) (EC 3.6.4.13) (ATP-dependent RNA helicase eIF4A-2) | |
| **P35700** | P35700 | PRDX1_MOUSE | Peroxiredoxin-1 (EC 1.11.1.15) (Macrophage 23 kDa stress protein) (Osteoblast-specific factor 3) (OSF-3) (Thioredoxin peroxidase 2) (Thioredoxin-dependent peroxide reductase 2) | Prdx1 |
| **Q9CQM5** | Q9CQM5 | TXD17_MOUSE | Thioredoxin domain-containing protein 17 (14 kDa thioredoxin-related protein) (TRP14) (Protein 42-9-9) (Thioredoxin-like protein 5) | Txndc17 |
| **E9PZJ2** | E9PZJ2 | E9PZJ2_MOUSE | Interferon regulatory factor 9 | Irf9 |
| **Q91XB0** | Q91XB0 | TREX1_MOUSE | Three-prime repair exonuclease 1 (EC 3.1.11.2) (3'-5' exonuclease TREX1) (DNase III) | Trex1 |
| **Q05910** | Q05910 | ADAM8_MOUSE | Disintegrin and metalloproteinase domain-containing protein 8 (ADAM 8) (EC 3.4.24.-) (Cell surface antigen MS2) (Macrophage cysteine-rich glycoprotein) (CD antigen CD156a) | Adam8 |
| **Q5F2E7** | Q5F2E7 | NUFP2_MOUSE | Nuclear fragile X mental retardation-interacting protein 2 (82 kDa FMRP-interacting protein) (82-FIP) (FMRP-interacting protein 2) | Nufip2 |
| **Q3TC45** | Q3TC45 | Q3TC45_MOUSE | Protein S100-A10 (S100 calcium binding protein A10 (Calpactin), isoform CRA_a) (S100 calcium binding protein A10 (Calpactin), isoform CRA_b) | S100a10 |

| | | | | |
|---|---|---|---|---|
| **J3QNE8** | J3QNE8 | J3QNE8_MOUSE | Putative sodium-coupled neutral amino acid transporter 10 | Slc38a10 |
| **P48024** | P48024 | EIF1_MOUSE | Eukaryotic translation initiation factor 1 (eIF1) (Protein translation factor SUI1 homolog) | Eif1 |
| **Q3UPN9** | Q3UPN9 | Q3UPN9_MOUSE | Putative uncharacterized protein | Cebpb |
| **P22893** | P22893 | TTP_MOUSE | Tristetraprolin (TTP) (Growth factor-inducible nuclear protein NUP475) (Protein TIS11A) (TIS11) (TPA-induced sequence 11) (Zinc finger protein 36) (Zfp-36) | Zfp36 |
| **Q64339** | Q64339 | ISG15_MOUSE | Ubiquitin-like protein ISG15 (Interferon-induced 15 kDa protein) (Interferon-induced 17 kDa protein) (IP17) (Ubiquitin cross-reactive protein) | Isg15 |
| **Q8CBB9** | Q8CBB9 | RSAD2_MOUSE | Radical S-adenosyl methionine domain-containing protein 2 (Viperin) (Virus inhibitory protein, endoplasmic reticulum-associated, interferon-inducible) | Rsad2 |
| **Q8BGW5** | Q8BGW5 | SOSB2_MOUSE | SOSS complex subunit B2 (Nucleic acid-binding protein 1) (Oligonucleotide/oligosaccharide-binding fold-containing protein 2A) (Sensor of single-strand DNA complex subunit B2) (Sensor of ssDNA subunit B2) (SOSS-B2) (Single-stranded DNA-binding protein 2) | Nabp1 |
| **O09172** | O09172 | GSH0_MOUSE | Glutamate--cysteine ligase regulatory subunit (GCS light chain) (Gamma-ECS regulatory subunit) (Gamma-glutamylcysteine synthetase regulatory subunit) (Glutamate--cysteine ligase modifier subunit) | Gclm |
| **Q9EQU5-2** | Q9EQU5-2 | SET_MOUSE | Protein SET (Phosphatase 2A inhibitor I2PP2A) (I-2PP2A) (Template-activating factor I) (TAF-I) | |
| **Q4FJM5** | Q4FJM5 | Q4FJM5_MOUSE | Ras homolog gene family, member B (Rho-related GTP-binding protein RhoB) (Rhob protein) | Rhob |
| **Q3U472** | Q3U472 | Q3U472_MOUSE | CD274 antigen (PD-L1) (Programmed cell death 1 ligand 1) | Cd274 |
