## Supplemental Table 3 for "Channeling macrophage polarization via selective translation inhibition by rocaglates increases macrophage resistance to Mycobacterium tuberculosis"

**Supplementary Table 3.**

| Property | Rocaglate | IFNγ | IFNγ references |
|---|---|---|---|
| **STAT1 activation** | - | + | |
| **Translation Inhibition** | + | + | **(*1*)** |
| **P38 MAPK activation** | + | + | **(*2*)** |
| **Autophagy Induction** | + | + | **(*3-6*)** |
| **IL4 suppression** | + | + | **(*7-9*)** |

**References:**


1. X. Su *et al.*, Interferon-gamma regulates cellular metabolism and mRNA translation to potentiate macrophage activation. *Nat Immunol* **16**, 838-849 (2015).
2. T. Matsuzawa *et al.*, IFN-gamma elicits macrophage autophagy via the p38 MAPK signaling pathway. *J Immunol* **189**, 813-818 (2012).
3. M. G. Gutierrez *et al.*, Autophagy is a defense mechanism inhibiting BCG and Mycobacterium tuberculosis survival in infected macrophages. *Cell* **119**, 753-766 (2004).
4. J. D. MacMicking, G. A. Taylor, J. D. McKinney, Immune control of tuberculosis by IFN-gamma-inducible LRG-47. *Science* **302**, 654-659 (2003).
5. A. R. Shenoy *et al.*, Emerging themes in IFN-gamma-induced macrophage immunity by the p47 and p65 GTPase families. *Immunobiology* **212**, 771-784 (2007).
6. S. B. Singh, A. S. Davis, G. A. Taylor, V. Deretic, Human IRGM induces autophagy to eliminate intracellular mycobacteria. *Science* **313**, 1438-1441 (2006).
7. H. L. Dickensheets, R. P. Donnelly, Inhibition of IL-4-inducible gene expression in human monocytes by type I and type II interferons. *J Leukoc Biol* **65**, 307-312 (1999).
8. T. Naka *et al.*, SOCS-1/SSI-1-deficient NKT cells participate in severe hepatitis through dysregulated cross-talk inhibition of IFN-gamma and IL-4 signaling in vivo. *Immunity* **14**, 535-545 (2001).
9. C. R. Yu *et al.*, Cell proliferation and STAT6 pathways are negatively regulated in T cells by STAT1 and suppressors of cytokine signaling. *J Immunol* **173**, 737-746 (2004).
